# The visuomotor transformations underlying target-directed behavior

**DOI:** 10.1101/2024.05.07.592863

**Authors:** Peixiong Zhao, Yuxin Tong, Ivan P. Lazarte, Biswadeep Khan, Guangnan Tian, Kenny K. Y. Chen, Thomas K. C. Lam, Yu Hu, Julie L. Semmelhack

## Abstract

The visual system can process diverse stimuli and make the decision to execute appropriate behaviors, but it remains unclear where and how this transformation takes place. We imaged the zebrafish visual system while larvae responded with hunting, freezing, and escape behaviors, and systematically identified visually driven neurons and behaviorally correlated sensorimotor neurons. Our analyses indicate within the optic tectum, broadly tuned sensory neurons are functionally connected to sensorimotor neurons that respond specifically during one behavior, transforming visual information into motor output. We also identified sensorimotor neurons in four other areas downstream of the tectum, and these neurons are also specific for one behavior, indicating that once the decision to behave has been made, the segregation of the pathways continues in later areas. Our findings suggest that the tectum receives visual sensory information and is responsible for selecting a single behavioral outcome, which is then relayed to downstream areas.

**Significance statement:** Here, we developed a novel visually-evoked freezing paradigm in zebrafish, and combined this with escape and hunting behaviors to ask how visual stimuli are identified and converted into different behavioral outcomes. We found that the optic tectum contains neurons that detect all three stimuli, as well as sensorimotor neurons for the three behaviors, suggesting that it is a site of sensorimotor transformation, which was supported by our analysis of correlations between the populations. The sensorimotor neurons in the tectum are highly specific for one behavior, and this segregation is maintained in the three downstream areas where we also identified sensorimotor neurons, indicating that the tectum flexibly transforms visual information into a single behavioral output.

## Introduction

Animals use their visual system to detect an array of visual targets, and respond with appropriate behaviors. Prey stimuli trigger orientation, approach, and consumption behaviors^1–3^, while predator-like stimuli can evoke fighting, fleeing or freezing^4,5^. Fleeing involves rapid movement, which may be directed away from the potential threat^6–9^. Freezing, on the other hand, is a cessation of movement, which may help the animal avoid detection, accompanied by a decrease in heart rate, or bradycardia^10–13^.

In many systems, these behaviors can be evoked by simple visual stimuli; small moving objects represent prey and trigger hunting behavior^14,15^, while looming stimuli can represent an approaching predator and trigger escape^7–9,16^. In contrast, an object sweeping across the visual field without changing size has been shown to evoke freezing in both flies and mice^17–19^, and this stimulus may mimic a predator that is cruising but not approaching. The detection of these stimuli within the early visual system is partially understood^7,8,15,20,21^, but it remains unclear, particularly in vertebrates, how the visual information is transformed into the appropriate behavioral output.

In order to perform these visual behaviors, the animal must (1) detect the identity and position of the stimulus, (2) decide whether and how to respond, and (3) execute a directed motor program toward or away from the target (in the cases of hunting and escape). The decision of whether to respond and with which behavior may be dependent on satiety, context, locomotor state, or other kinds of sensory input^22–25^. For target-directed behaviors, which are triggered by visual stimuli located at a particular point in space, the optic tectum (superior colliculus in mammals) is a likely site for visuomotor transformation, as it receives retinal input, contains a map of visual space, has outputs to many motor areas, and receives a variety of other sensory and state inputs^26–28^. However, other brain areas have also been implicated in these behaviors^29–31^, and their manipulation has been shown to affect hunting and defensive behaviors^32–34^. In addition, it has been difficult to comprehensively record from these areas, particularly during behavior, in order to observe how the animal decides whether to respond, and selects the appropriate type of behavior.

Here, we first characterize visually evoked freezing (immobility and bradycardia) in response to a sweeping stimulus in zebrafish larvae. In addition to flies^23^, rodents^17^, and humans^11^, freezing behavior has also been found in zebrafish; adult zebrafish exhibit immobility in response to threatening stimuli such as novel environments or alarm pheromone^35,36^, and late-stage larvae have a bradycardia response during conditioned fear^12,37^. However, visually evoked freezing had not been described in zebrafish. To investigate the sensorimotor circuits mediating this behavior, and how they relate to the pathways for other visual behaviors, we presented the sweeping stimulus as well as prey and looming stimuli, and observed whether the larvae responded by freezing, hunting, or escaping. We used volumetric 2p calcium imaging to map the sensory neurons that respond to predator-like and prey stimuli, and the sensorimotor neurons that are correlated with stimulus-evoked freezing, escape, and hunting. We identified sensory neurons for the stimuli in the tectum, and found that sensorimotor neurons were also located there, as well as in several other downstream areas. Our data suggest that the tectum is a site of a visuomotor transformation that converts relatively broad sensory input into segregated pathways for each behavior.

## Results

### A large translating disk causes immobility

To test whether a visually evoked freezing response could be triggered in zebrafish larvae, we designed a 15° disk stimulus that would sweep horizontally across the frontal visual field. This initial stimulus was similar in size to that used to trigger freezing in other systems^17,19^, and larger than a prey stimulus^15^. We projected the dark sweeping disk on a red background onto the screen of our behavior chamber (figure 1A). Within the chamber, a head-fixed 6- or 7-day post fertilization larva was mounted 1 cm from the screen, and we used high speed cameras above and to the side to monitor its behavior (figure 1A, insets). The larvae would intermittently perform spontaneous swims, and we found that the presentation of the sweeping stimulus would suppress swimming for several seconds (Supplementary video 1). When we quantified swim probability (probability of a swim bout occurring in that frame) across all sweeping stimulus trials from several larvae, we found that there was a decrease in swim probability from stimulus onset for a period of about 10 seconds (S Supplementary figure 1A, upper panel), This decrease can be quantified as a change in swim probability that was significantly greater for the stimulus vs. no stimulus trials (Supplementary figure 1B). This suggests that the sweeping stimulus was suppressing spontaneous swims and causing immobility, one of the signatures of freezing.

**Figure 1:**
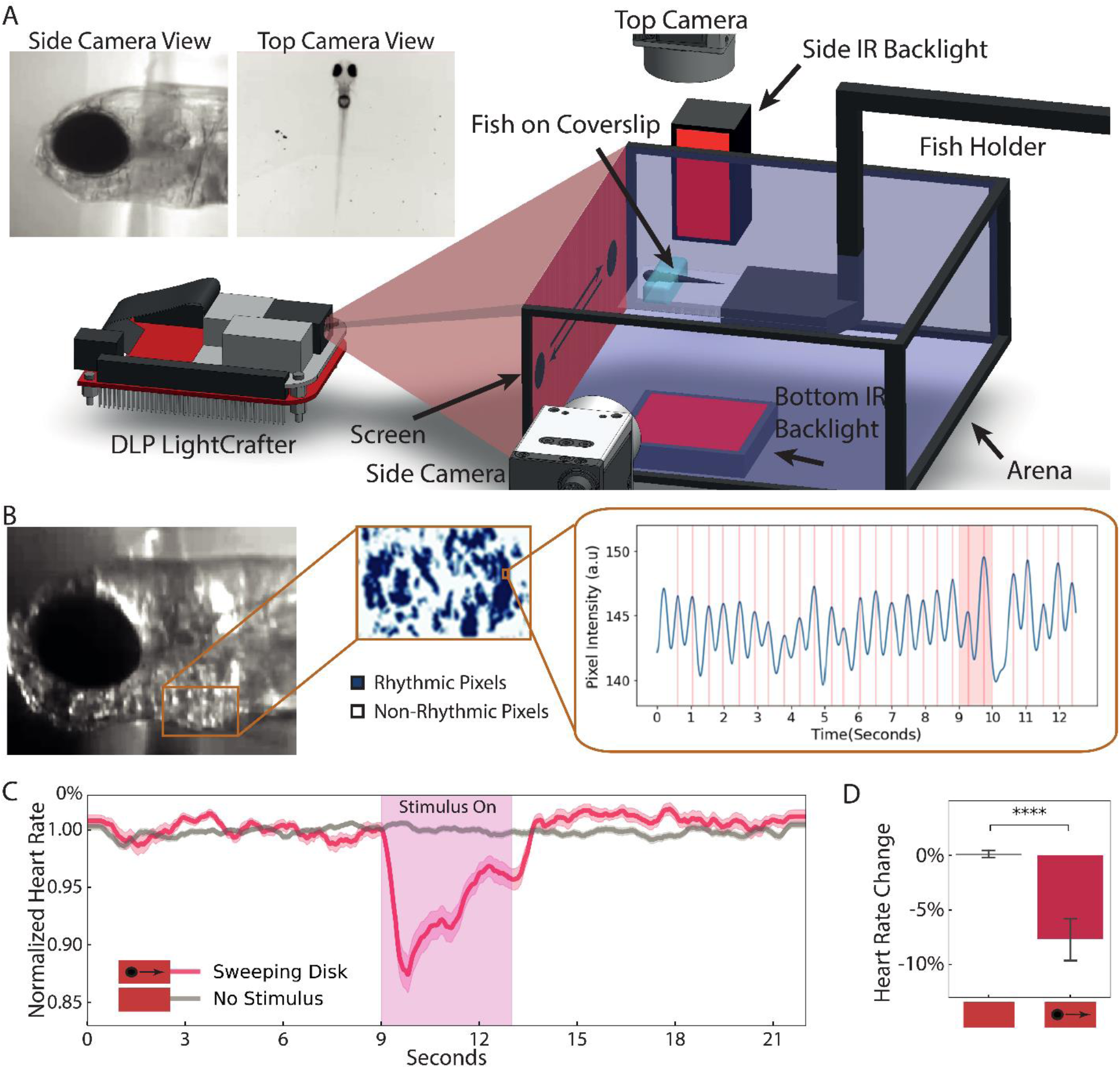
A large translating disk causes immobility and bradycardia. (A) Schematic of the experimental setup. (B) Identification of rhythmic pixels within the heart ROI, and pixel intensity peaks corresponding to heart beats. (C) Normalized heart rate in response to sweep or no stimulus. Shading indicates Standard Error. (D) Change in heart rate during the 3 seconds after stimulus onset. n = 14 larvae. Error bar = Standard Deviation. ****, p<0.0001, Mann–Whitney U test.

### Bradycardia can be used as an indicator of freezing

In many species, freezing is accompanied by a decrease in heart rate that occurs at the onset of immobility^10,11,38^. Therefore, we asked whether our putative freezing stimulus would also cause bradycardia, in addition to suppressing movement. We used the side camera to record the heart beating, and analyzed heart rate similarly to Mann et. al.^13^ by measuring the change in intensity of rhythmically active pixels (methods). We calculated the heart rate for each frame of the video by finding the distance between local peaks in the intensity curve of the rhythmically active pixels (figure 1B). We found that the dark sweeping disk robustly triggered a reduction in heart rate at the onset of stimulus presentation (figure 1C, and Supplementary video 2), and the heart rate change during the stimulus time window was significantly greater than the no stimulus case (figure 1D). We next asked whether bradycardia and the decrease in movement were linked, by plotting the swim probability in bradycardia vs. non-bradycardia trials. For this analysis, we defined bradycardia as a decrease in heart rate to ceiling greater three standard deviations from the mean (methods) starting during the 9-13 second window, in either stimulus or no stimulus trials. We found that bradycardia was associated with a decrease in swimming probability during and for several seconds after the stimulus (Supplementary figure 1A, lower panel), although the stimulus did evoke short-latency movement in a small minority of trials. Overall, trials with bradycardia had significantly less movement than those without (Supplementary figure 1C). Taken together, these data suggest that, as in other systems, a dark sweeping visual stimulus of constant size triggers the innate freezing response to potential predators, which can be observed as a cessation of movement and simultaneous decrease in heart rate. As spontaneous swim probability is quite variable between larvae, we used bradycardia as the behavioral readout for subsequent experiments.

### Optimal size and speed parameters to induce bradycardia

We next explored the stimulus parameters to identify those that would most effectively induce bradycardia. We presented dark sweeping disks on red, UV, and green backgrounds, and found that all were effective at triggering bradycardia (Supplementary figure 1D), and the heart rate change during the stimulus window was significantly larger than for the no stimulus condition (Supplementary figure 1E). In comparison, a bright green disk on a dark background evoked some decrease in heart rate, but was less effective than the dark stimuli (Supplementary figure 1D and E), and UV and red sweeping disks on a dark background similarly caused only a small decrease in heart rate (Supplementary figure 1F and G). We therefore chose to use the dark disk on a red background for further experiments, due to its greater with 2p imaging. Next, we varied stimulus speed, and found that a 15° diameter disk moving with a broad range of speeds effectively triggered bradycardia (Supplementary figure 1H), and we chose 60°/second as a value in the middle of that range. Using a speed of 60°/second, we found that stimuli of 5-30° degrees in diameter strongly reduced the heart rate (Supplementary figure 1I), and we chose to use a 15° diameter disk moving at 60°/second for further experiments.

### Functional imaging to identify sensory neurons that respond to each stimulus

To investigate how larvae detect the sweeping visual stimulus and respond with freezing behavior, we adapted our behavioral assay for volumetric 2p calcium imaging. We chose two other behaviorally relevant visual stimuli to contrast with the sweep stimulus. The first was a prey stimulus, a small (4° diameter) sweeping UV dot that has a similar horizontal trajectory and speed as the sweep stimulus, but nonetheless triggers hunting behavior (Supplementary video 3, Supplementary figure 2A)^39,40^, as defined by eye convergence and prey capture swims^15,41^. The second was a looming stimulus, which is, like the sweeping disk, a dark object, but expanding from 6 to 60° in diameter (Supplementary figure 2A). This stimulus primarily evokes escape (Supplementary video 4), which involves a high velocity and amplitude turn away from the stimulus^7,8^. We presented each stimulus eight times, with a two-minute interstimulus interval. We used larvae expressing nuclear localized GCaMP6s pan-neuronally (*elavl3:Hsa.H2B-GCaMP6s*^42^, aka *elavl3*) and an electrically tunable lens to image a large volume of the brain, including many of the areas thought to be involved in defensive behaviors (figure 2A). We recorded from 14 planes at 2 Hz, covering about 2/3 of the brain by volume (figure 2A, Supplementary table 1). We used suite2p^43^ to motion correct and segment cell bodies, identifying a total of 188,272 cell bodies within our dataset of 7 fish, and mapped the cell bodies onto the mapZebrain atlas^44^. Within the total population of segmented cell bodies, we then defined a population of active neurons (correlated with one of the stimuli or behaviors in 30% of the relevant trials (methods): 63,420 neurons in total) to streamline the subsequent analysis.

We first identified the purely stimulus-correlated neurons, which we call “sensory neurons”, for each stimulus. We adopted an approach similar to that of a recent study^45^, based on the idea that sensory neurons should respond to repeated presentation of the same stimulus with high periodicity. For each neuron, we calculated its average response to a given stimulus over eight repeats, and generated the average trace (Trace_avg_) by concatenating the average response eight times. We then calculated a Sensory Index (SI) for the neuron’s response to that stimulus, which reflects the variance explained by the periodic component of the response (methods), such that if the average and actual trace have the same variance, the SI will have the maximum value (figure 2B). Neurons with an SI in the first percentile of the population had high periodicity, responding every time the stimulus was presented, while neurons with lower SI were less regular in their responses (Supplementary figure 2B).

**Figure 2:**
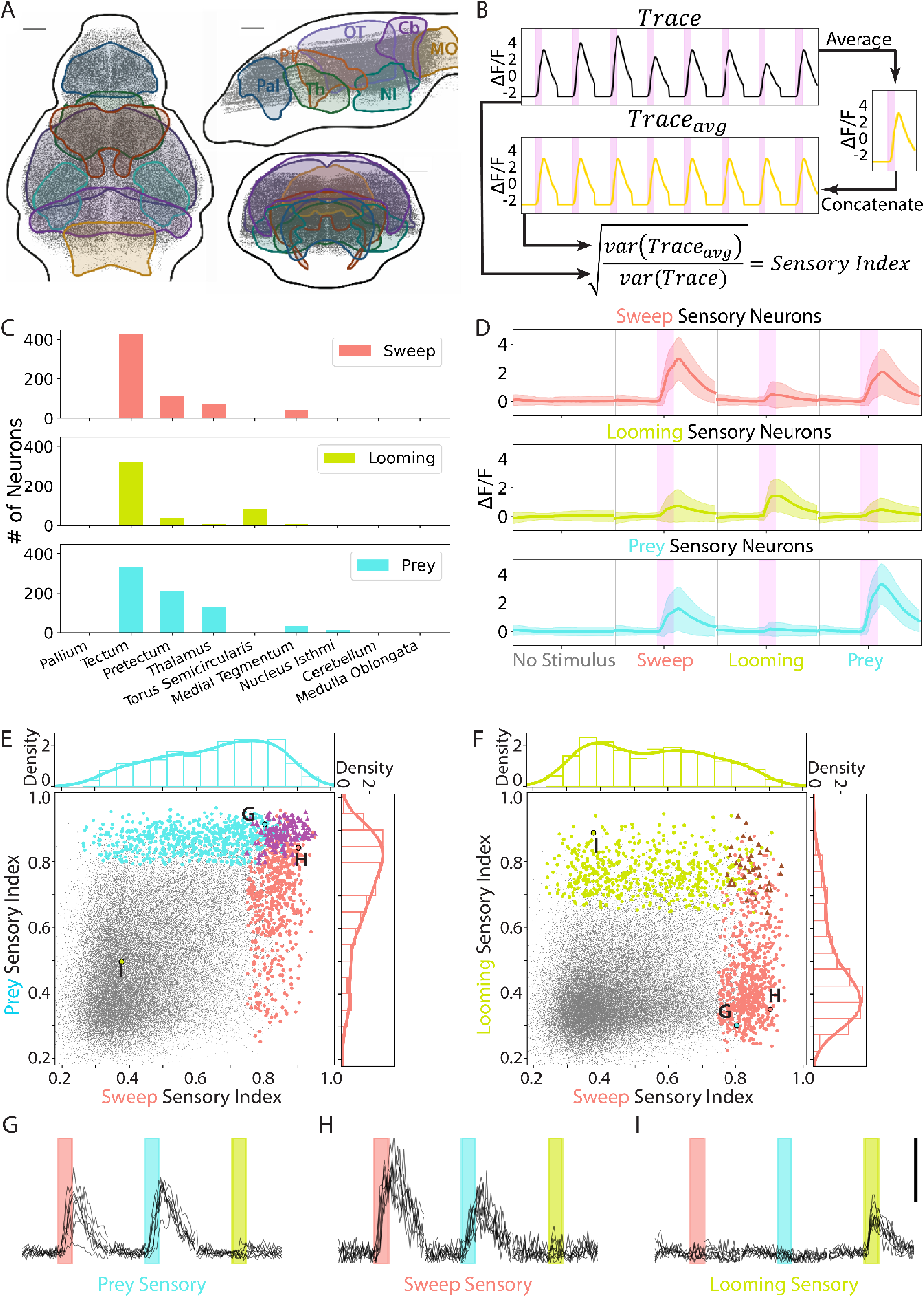
Identification of sensory neurons for sweep, prey and looming stimuli. (A) Cell bodies in the imaging dataset (n = 7 larvae, 188,272 neurons). Pallium, Pal; Pretectum, Pt; Thamalus, Th; Optic Tectum, OT; Nucleus Isthmi, NI; Cerebellum, Cb; Medulla Oblongata, MO. Scale bar = 50 µm. (B) Calculation of the sensory index (SI) of each neuron. (C) Number of sensory neurons in each brain area. (D) The average response of each sensory neuron population to all stimuli. Pink bars = 4-second stimulus display. Shading indicates standard deviation. (E) SI of all sweep (red), prey (blue), and prey + sweep (purple) sensory neurons, and all active neurons in the dataset (grey, 63,420 neurons). Density plots represent sensory neurons of each type. (F) SI of looming (green), sweep (red), and looming + sweep (brown) sensory neurons. (G-I) Response of example sweep, prey, and looming sensory neurons selected near the mode (i.e., max density) of that population’s density plot. Red, blue and green bars indicate 4-second sweep, prey or looming stimuli. Scalebar indicates ΔF/F = 3.

Use of periodicity alone to identify sensory neurons gave us a significant proportion of neurons with flat or slowly ramping traces. Therefore, to identify sensory neurons for each stimulus, we first selected neurons that were correlated with the stimulus regressor (top 10%, see methods), then within that population took the top 15% based on the SI. Neurons at the lower limit of this range still responded very regularly to the stimulus (Supplementary figure 2B, middle trace), so we chose 15% as a threshold that would allow us to identify a relatively large population of sensory neurons, while still ensuring that whole population is highly periodic. We further filtered the population for neurons that were colocalized across multiple animals (spatial colocalization test, methods)^46^. Within our population of active neurons, we identified 682 sweep, 556 looming, and 746 prey sensory neurons in total. We found that the vast majority of the sensory neurons for the all three stimuli were in the optic tectum, while the prey stimulus also activated a large population within the pretectum (figure 2C), consistent with previous studies^15,47^. To validate our selection of 15% of the SI as a threshold, we compared the anatomical distribution of sensory neurons for thresholds ranging from 5% to 30% SI, and found that the fraction of neurons in each brain area was consistent (Supplementary figure 2C-E), suggesting that no major populations of sensory neurons are excluded by our chosen threshold. We also plotted the proportion of sensory neurons from each larva, and found that these reliably stimulus-driven neurons in the tectum were present in all seven fish (Supplementary figure 2F).

We next examined the functional properties of the three populations of sensory neurons, and found that sweep sensory neurons on average responded robustly to the sweeping disk, as expected, and interestingly, they also responded to the prey dot (figure 2D, red). Looming sensory neurons responded primarily to the looming stimulus (figure 2D, green), while prey sensory neurons on average responded to the sweeping disk as well as to the prey dot (figure 2D, blue). To further examine the functional properties across stimuli at the individual neuron level, we use a scatter plot to visualize each neuron’s sweep sensory index (SI) vs. its prey SI. As expected, most sweep sensory neurons had a high sweep SI (figure 2E, red dots), and a large proportion of them also had a prey SI of ∼0.8 (figure 2E, red density plot). These values reflect a neuron that responded with high periodicity to the sweep stimulus, and nearly as regularly to the prey stimulus. Indeed, the response of an example neuron from the mode of the sweep sensory neuron distribution shows a fairly periodic response to the prey stimulus (figure 2H). Similarly, the prey sensory neurons (figure 2E, blue) all had a high prey SI, and many of them also had a sweep SI of ∼0.8 (figure 2E, blue density plot), indicating regular activation by the sweep stimulus (figure 2G). There was also a significant population of neurons that were classified as both sweep and prey sensory neurons (figure 2E, purple triangles). In contrast, most sweep sensory neurons had a relatively low looming SI of ∼0.4 (figure 2F, red density plot), indicating a lack of response to the looming stimulus (figure 2H). Looming sensory neurons similarly had low SI for sweep (figure 2F, light green density plot, and figure 2I). These results suggest that the tectal sensory neurons may be encoding object location more specifically than object identity (i.e. prey vs. predator), as there was substantial overlap between the stimuli with similar trajectories (sweep and prey), but not between the two dark, predator-like stimuli (sweep and looming).

### Identification of sensorimotor neurons correlated with each behavior

Given that animals do not respond with the identical behavior to each repeat of a visual stimulus, at some point within the visuomotor pathway visual information must be integrated with state, history, or other sensory information to make the decision to behave. The neurons at this point and later in the pathway can be thought of as “sensorimotor” neurons (SM neurons), and will be correlated with a behavior which is evoked by a particular visual stimulus. To search for these neurons in our imaging dataset, we first classified the behaviors triggered by the three stimuli as hunting, freezing, and escapes. Trials containing eye convergence bouts during stimulus presentation were classified as hunting, those with fast, large amplitude swims were classified as escape, those with no movement and bradycardia were classified as freezing, and those with multiple types of behavior during the stimulus window were excluded from the analysis (Supplementary figure 3A-B). Based on this classification, the sweeping stimulus triggered mostly freezing and some escape, while the looming stimulus largely evoked escape and a smaller percentage of freezing trials (figure 3A).

**Figure 3:**
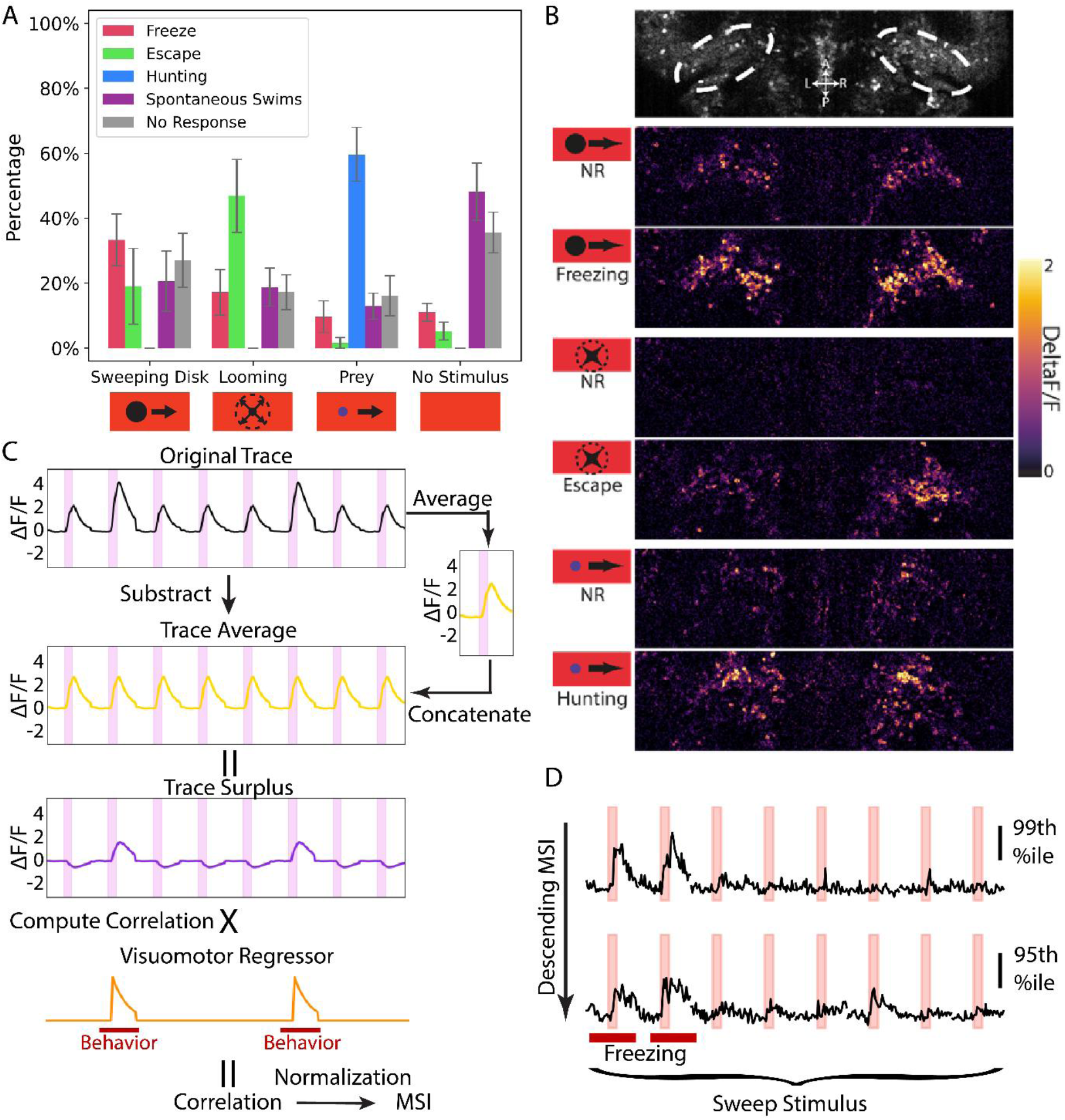
Identification of sensorimotor (SM) neurons for freezing, escape, and hunting. (A) Percent of trials with each behavioral response to the stimuli. Error bar represents Standard Deviation. n = 7 larvae (B) Example responses of cell bodies in the nucleus isthmi (NI) to sweep, looming, and prey stimuli in no behavioral response (NR) and freezing, escape, and hunting trials. (C) Schematic calulation of freezing MSI for a neuron in a fish with freezing in the second and 6^th^ trials. First, the trace average is subtracted from the origingal trace to generate the trace surplus, then the Pearson correlation between the trace surplus and the visuomotor regressor is computed and normalized to give the freezing motor surplus index (MSI) for that neuron. (D) Example ΔF/F neuronal traces during the eight sweep presentations from a larva with two freezing trials. Neurons are from the 99^th^ and 95^th^ percentile of the freezing MSI. Red bars indicate freezing trials.

Within certain brain areas, we observed that some neurons had little response to our stimuli in trials without behavior, but were robustly activated in behavior trials, with different sets of neurons responding during the three types of behaviors (nucleus isthmi, figure 3B). To systematically identify these SM neurons, again inspired by the same prior study^45^, we designed an analysis method that would decompose neuronal responses into stimulus-dependent and - independent components, to allow us to identify a population of SM neurons that would be truly behavior- and not stimulus-correlated. This approach avoids the issues of using a simple regression analysis for data where the stimuli and behaviors are correlated, which can result in significant overlap between two populations. For each neuron, we calculated a “trace surplus”, or stimulus independent trace, by subtracting the average response to a given sensory stimulus. We then calculated the Pearson correlation between the trace surplus and a visuomotor regressor (methods, Supplementary figure 4), and used that correlation score to calculate a Motor Surplus Index (MSI) for the primary behavior driven by each stimulus (figure 3C). To evenly distribute the SM neurons among animals, we normalized each fish’s MSI scores such that 0 was the minimum score, and 1 was the maximum score.

As expected, neurons with a high MSI responded more robustly in behavior trials, whereas lower scoring neurons were less correlated with behavior (figure 3D. To identify three populations of SM neurons for further analysis, we had to select a threshold within the MSI values. To determine an appropriate MSI threshold, we calculated the motor enhancement, or difference between no response and behavior trials of the sensorimotor neuron populations. For example, for freezing sensorimotor neurons, the motor enhancement is the difference between the peak ΔF/F in freezing trials to the peak ΔF/F in no response trials of the sweep stimulus (Supplementary figure 5A, red line). Motor enhancement was greatest in the 99^th^ percentile neurons by MSI, and lower but still present in the 90^th^ percentile (Supplementary figure 5B). We found that there was an elbow in the degree of motor enhancement around the 97^th^ percentile for all behaviors (Supplementary figure 5C), meaning that selecting the top 3% of neurons allows us to capture the neurons most modulated by the behaviors. We thus defined the 97^th^ percentile of freezing MSI as the threshold for freezing SM neurons, and for each fish classified the neurons with MSI above this threshold as SM neurons, and the same for hunting and escape. After spatial colocalization test, we were left with 1219 freezing, 1239 escape, and 1262 hunting SM neurons in total.

Using this approach, we were able to identify populations of SM neurons that had on average a more robust response during behavior trials, in contrast to their sensory neuron counterparts (figure 4A-C). One surprising finding was that each set of SM neurons had a barely perceptible average response to their sensory stimulus during no behavior trials; i.e., freezing SM neurons had virtually no response to the sweep stimulus in trials without freezing, and escape and hunting SM neurons behaved similarly (figure 4A-C, NR trials). This low responsiveness to sensory stimuli was not due to the 97^th^ percentile threshold we used for SM neurons; the average response in no behavior trials was consistently low for the 99^th^ to 70^th^ percentile of neurons by MSI score (Supplementary figure 5C, grey lines).

**Figure 4:**
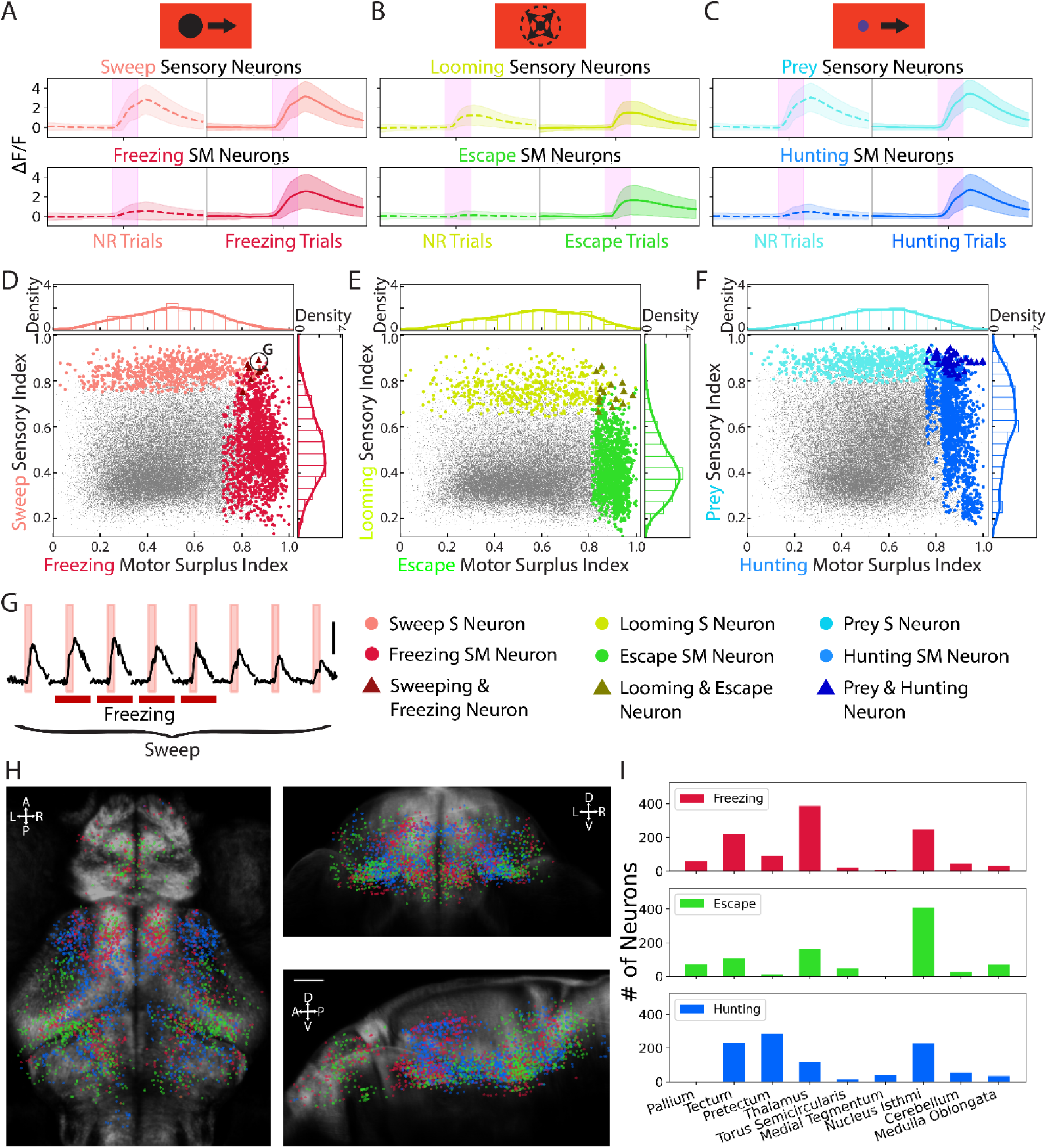
Functional properties and anatomical locations of sensorimotor (SM) neurons for freezing, escape, and hunting. (A-C) Average calcium response of the sensory and SM neurons during behavior and no response trials. (D) The distribution of sweep sensory neurons (light red) and freezing SM neurons (dark red) on the sweep SI and freezing MSI. Crimson triangles: neurons belonging to both populations. (E) The distribution of the looming sensory neurons (light green) and escape SM neurons (dark green) on the looming SI and escape MSI. Grey triangles: neurons belonging to both populations. (F) The distribution of the prey sensory neurons (light blue) and hunting SM neurons (dark blue) on the prey SI and hunting MSI. Navy triangles: neurons belonging to both populations. (G) Response of example neuron with both high sweeping SI and freezing MSI, annotated in D. Light red bars represent 4-second presentation of sweep stimuli. Dark red bars represent trials with a freezing response. Scale bar indicates ΔF/F = 3. (H) Locations of SM neurons in the recording volume. Red: Freezing neurons. Green: Escape neurons. Blue: Hunting neurons. Scale bar is 50 µm. (I) Numbers of SM neurons in each brain area.

To look more closely at the response properties of individual neurons within the SM populations, we plotted SI vs. MSI of each stimulus and behavior pair. We found that the freezing SM neurons (figure 4D, dark red) typically have a low sweep SI, with the mode of the distribution around 0.4, indicating little periodicity for the sweep stimulus. Although it is possible for neurons to be classified as both sweep sensory and freezing SM neurons (maroon triangles), as long as they respond periodically to the stimulus and have some motor surplus in the behavior trials, there were only eight neurons of this type in our entire imaging volume (figure 4D and example trace, figure 4G). The counts of looming + escape (figure 4E) and prey + hunting populations (figure 4F) were similarly low. Thus, this analysis allows us to identify populations of neurons that are correlated with the three behaviors triggered by the visual stimuli. These SM neurons typically have very little response to the corresponding stimulus itself, and therefore there was virtually no overlap between sensory and SM populations.

We next looked at the anatomical location of each SM neuron type within the mapZebrain atlas^44^. We found that SM neurons for the three behaviors were distributed in distinct patterns within our imaging volume (figure 4H). There was a significant population of hunting, freezing and escape SM neurons in the tectum, as well as in the thalamus and nucleus isthmi (NI) (figure 4I). A large population of hunting SM neurons was also found in the pretectum (figure 4I), consistent with previous findings in zebrafish^32^. The anatomical locations of SM neurons were fairly consistent across fish, with each larva having some SM neurons in the major areas (Supplementary figure 5D). We also compared the locations of the top 600 neurons in each fish by MSI prior to within fish normalization and the spatial colocalization test, and found that the patterns and brain regions were still similar across fish (Supplementary figures 6-8), although there were subtle differences, for example in the number of freezing SM neurons in the thalamus. This may be due to differing numbers of behavior trials between fish; in a fish with several freezing trials (e.g. fish 6, Supplementary figure 6A), many neurons in the thalamus have a high MSI, and these will take up a larger share of the top 600. In contrast, a fish with only one freezing trial, e.g. fish 1, will have more randomly distributed neurons, many of which will be eliminated by the spatial colocalization test.

To further validate our method of selecting SM neurons, we next asked whether the activity of an SM population can predict behavior. We conducted a separate selection of freezing SM neurons, using the freezing annotation and neuronal activity of seven trials, with one trial held out (Supplementary figure 9A). We then compared the held-out trial activity to the maximum population activity during the freezing and no response trials (Supplementary figure 9B), to ask if they could predict whether the held-out trial was a freezing or no response trial. We also asked how well a set of null SM neurons, selected based on a permuted behavioral annotation (Supplementary figure 9C) could predict the permuted annotation, and found that the real freezing SM neurons were significantly better than the null population at predicting whether the larva froze in the held-out trial, with an accuracy of 84.8% (Supplementary figure 9D), although fish with only one freezing trial (fish 4 and 5, Supplementary figure 10) or trials removed due to escape responses (fish 1, Supplementary figure 10) may have had insufficient data to allow an accurate prediction. In summary, our method of selecting SM neurons identified populations of neurons that responded robustly during behavior trials, were separate from the corresponding sensory populations, were located in consistent regions across fish, and could predict behavioral responses.

### Sensorimotor transformation and pathway divergence within the tectum

In addition to containing the three types of sensory neurons, we found that the optic tectum also contained SM neurons for all three behaviors (figure 5A). As the tectum was the area with by far the largest concentration of sensory neurons of the three types (figure 2C), and it also contains SM neurons for the three behaviors, it seems likely that the visual stimuli are detected in this area, and the decision to respond with freezing, escape or hunting could be made here, perhaps based on other inputs to the tectum. We therefore looked more closely at the anatomical distributions of SM neurons for clues as to how such transformations could occur.

**Figure 5:**
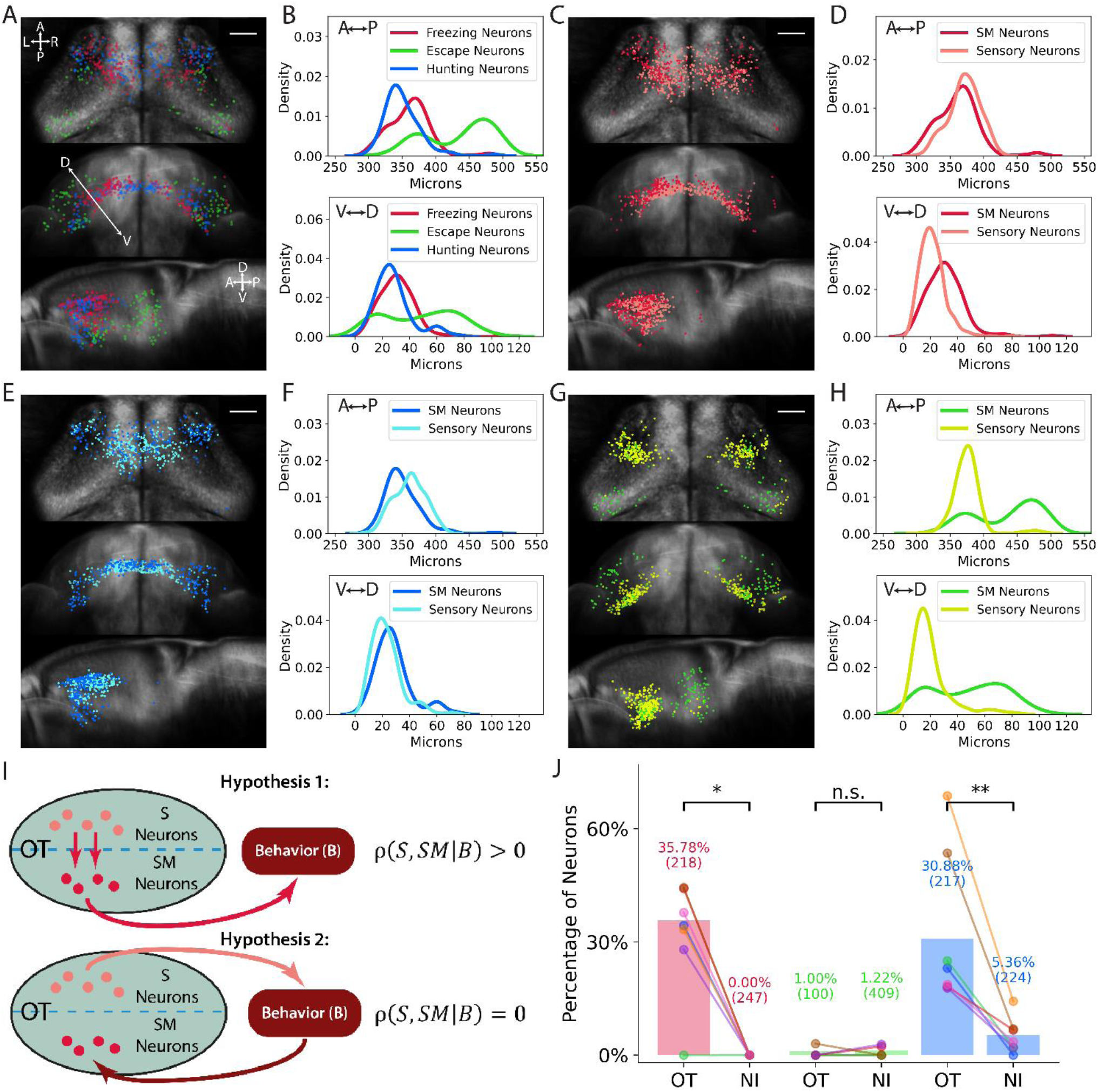
Sensory and sensorimotor neurons in the tectum. (A) Locations of the three types of SM neurons in the tectum. Scale bar = 50 µm. (B) Locations of SM neurons on the anterior/posterior and dorsal/ventral axes. (C-D) Locations of sweep sensory neurons (light red) and freezing SM neurons (dark red) in the tectum and their locations on the anterior/posterior and dorsal/ventral axes. (E-F) Locations of prey sensory neurons (light blue) and hunting SM neurons (dark blue) in the tectum and on the anterior/posterior and dorsal/ventral axes. (G-H) Locations of looming sensory neurons (light green) and escape SM neurons (green) in the tectum and their locations on the anterior/posterior and dorsal/ventral axes. (I) Hypotheses for the tectum S and SM connectivity in the sensorimotor circuit. (J) Percentage of SM neurons that were significantly correleated with the tectal sensory neurons. Colored dots respresent values for individual fish. In parenthesis is the total number of SM neurons of that type across all fish. NI = Percentage of SM neurons in the NI that were significantly correlated with tectal sensory neuron In all behaviors. Percentage-by-chance was 0% for both tectal and NI SM neurons (methods). *, p<0.05. **, p<0.01, One-sided Wilcoxon signed-rank test. The p-values were 0.011 for freezing, 0.787 for escape, and 0.007 for hunting.

Hunting, freezing, and escape SM neurons were located in distinct regions of the tectum (figure 5A), with hunting neurons in the most anterior position, freezing neurons in the middle, and escape neurons most posterior (figure 5B), consistent with previous results implicating anterior tectum in approach and posterior in avoidance behaviors^48,49^. To ask whether optogenetic stimulation of different regions of the tectum could produce our behaviors, we expressed ChR2 under control of *Gal4s:1013t*^50^, and used an optic fiber to activate the anterior vs. posterior tectum. We observed significantly more eye convergence bouts when we stimulated the anterior vs. posterior tectum (Supplementary figure 11). Stimulation of the posterior tectum produced more escape bouts than anterior tectal stimulation, although the number of escapes was very small. These results suggest that activating hunting and escape SM neurons in the anterior and posterior regions of the tectum, respectively, could mediate hunting and escape responses.

We next looked at where each set of sensory neurons was located relative to its sensorimotor pair. The sweep sensory and freezing SM neurons were located in similar regions of the tectum along the anterior/posterior axis (figure 5C-D), with the SM neurons in a slightly dorsal region to the sensory neurons. The prey sensory and hunting SM neurons were also located in similar regions of the tectum, with the sensory population slightly more posterior (figure 5E-F). In contrast, the anatomical separation between the looming sensory and escape SM neurons was more pronounced, with the escape population markedly more posterior and dorsal (figure 5G-H). These data suggest that the sensory and SM neurons of each pair constitute separate anatomical populations, which may be connected to each other.

### Causal inference analysis supports the tectum as a site of sensorimotor transformation

To ask whether sensory and SM neurons in the tectum are connected, we looked for correlations between the activities of sensory and SM neurons of each type. If sweep sensory neurons are connected to freezing SM neurons, we would expect the two populations to be correlated (figure 5I, hypothesis 1). However, it is also possible that the freezing SM neurons in the tectum are not receiving input from their sensory counterparts, but are nonetheless correlated to them due to indirect pathways via some other brain area that determines behavior (hypothesis 2). To distinguish the two scenarios, we adapted statistical methods from causal inference^51^, which are based on the partial correlation between the sensory and SM neurons while conditioning on the behavior *ρ*(*S*, *SM*|*B*) (figure 5I). Intuitively, only in hypothesis 1 are the sensory and SM neurons are still correlated once their relationship with the behavior is removed (i.e., conditioning on the behavior).

To remove the behavior-related activity, we calculated a residual activity trace for each SM neuron, analogous to the surplus trace in figure 3C. First, we compute behavior and non-behavior avergages by averaging the neuron’s trace across corresponding trials (Supplementary figure 12A, middle row). Both templates are then subtracted from the original trace to obtain the residual trace (Supplementary figure 12A, bottom row). The sensory neurons’ residuals are calculated in the same way. The partial correlation given behavior *ρ*(*S*, *SM*|*B*) is then calculated as the Pearson correlation coefficient between the SM and sensory residual. Next, we determined whether such partial correlations between the freezing SM and the sweep sensory neurons are significant. If the S and SM neurons are indeed connected, each SM neuron probably receives inputs from only a subset of sensory neurons. Therefore, for each SM neuron, we quantify its strongest correlations by calculating the 90^th^ percentile of its partial correlations with all sweep sensory neurons (Supplementary figure 12C, right). To check for significance, we compare this 90^th^ percentile against a null distribution where we perform exactly the same analysis on trial-shuffled data (permuted trace, Supplementary figure 12B), to obtain an empirical p-value (Supplementary figure 12C, D).

After adjustment for multiplicity by controlling the false discovery rate (FDR), we found that substantial fractions, 35.78% and 30.88%, of the freezing and hunting SM neurons have a significant partial correlation with corresponding sensory neurons (figure 5J). For escape SM neurons, we only observed a small fraction of 1.00% with a significant partial correlation with looming sensory neurons (however, this fraction is still well above the chance level of 0% caluclated from permuted SM neurons (methods)). According to causal inference theory, these results on partial correlations strongly support hypothesis 1, where a sensorimotor transformation occurs within the tectum via connections between S and SM neurons, at least for freezing and hunting behaviors. One caveat is that our analysis relies on partial correlations; thus, direct validation of the proposed circuit and causal effects, for example through perturbation experiments, is needed in future work.

To further validate our methods, we apply the causal inference analysis to a number of controls. First, if the tectal sensory neurons are uniquely providing input to tectal SM neurons, SM neurons in other areas such as the NI should not be as correlated with tectal sensory neurons. Using the same analysis pipeline, we found 0% and 5.36% of NI SM neurons for freezing and hunting were significantly correlated with tectal sensory neurons (figure 5J). Second, we verified that correlations between residuals are due to within-trial fluctuations in activity rather than the subtraction of the average traces (Supplementary figure 13). For each SM neuron, we permuted its trace across trials within behavior and non-behavior categories, which leaves the behavior average unchanged (Supplementary figure 13A and B). For these permuted traces, after subtraction of averages, the p-values for partial correlation significance were dramatically larger than real data (Supplementary figure 13C), and as a result only a tiny fraction of the SM neuron populations were significantly correlated under the same criterion (less than 0.1%, Supplementary figure 13D). Third, we confirm the observed partial correlation between tectal SM and sensory neurons are not due to artifacts related to two-photon scanning or imaging noise (Supplementary figure 14). We consider a surrogate population of sensory neuorns (S’ neurons) located close to each sensory neuron (average distance was 6, 11, and 7 µm for sweeping, looming, and prey pairs) and in the same imaging plane, but with average tectal SI values (Supplementary figure 14A). Applying the causal inference analysis to the S’ and SM neurons results in much lower fractions of significantly correlated SM neurons for freezing and hunting behaviors (Supplementary figure 14B). Taken together, these controls lend further support to our hypothesis that a sensorimotor transformation occurs within the tectum for freezing and hunting.

It is also possible that sensorimotor transformations could occur in the pretectum and thalamus, as these areas also contain sensory and sensorimotor neurons. We performed the partial correlation analysis for freezing and hunting behaviors in these two areas (escape could not be analyzed because only 8 escape SM neurons were found in the pretectum, and only 7 looming sensory neurons were found in the thalamus, across all fish.) We found that the fraction of SM neurons that were siginificantly correlated with sensory neurons within the same area were substantially lower for thalamus and pretectum than for the tectum (all less than 8%, Supplementary figure 15A), with the exception of pretectum hunting SM neurons, of which 24.62% were significantly correlated with pretectal prey sensory neurons. This suggests that the pretectal hunting SM neurons are integrating input from the nearby prey sensory neurons, which raises the question of why two different sensorimotor transformations would exist for prey stimuli. We hypothesized that a different kind of sensory input might be extracted in the pretectum and conveyed to the tectal hunting SM neurons. We found that indeed, the majority of hunting SM neurons in the pretectum had significant partial correlations with tectal hunting SM neurons, (Supplementary figure 15B), suggesting that hunting SM neurons in these two areas are connected.

### Tectal sensorimotor neurons respond specifically during one behavior

Although the existence of tectal SM neurons for these behaviors has been inferred based on stimulation experiments^27,48,49^, this dataset gives us the opportunity to assess their functional properties for the first time; for example, how do they respond to various visual stimuli, and during the different types of behavior? It could be that some of the tectal SM neurons are correlated with multiple behaviors, for example a neuron could respond in both freezing and escape trials and signal the presence of a predator, or during both freezing and hunting trials and encode the presence of a moving object. To test this, we looked at the responses of each class to the three behaviors, and found that on average the populations responded only during their primary behavior, for example, the freezing SM neuron population in the tectum responded robustly in sweep-induced freezing trials, but not in escape trials triggered by looming and minimally in hunting trials triggered by prey stimuli (red line, figure 6A). On an individual neuron level, there was only one tectal neuron that was classified as both an escape + freezing SM neuron and also three neurons that were classified as both a freezing + hunting SM neuron (figure 6B), which is surprising given the overlap of the prey and sweep sensory populations (figure 2E). These three “both” SM neurons represent 1.4% of the tectal freezing SM population.

**Figure 6:**
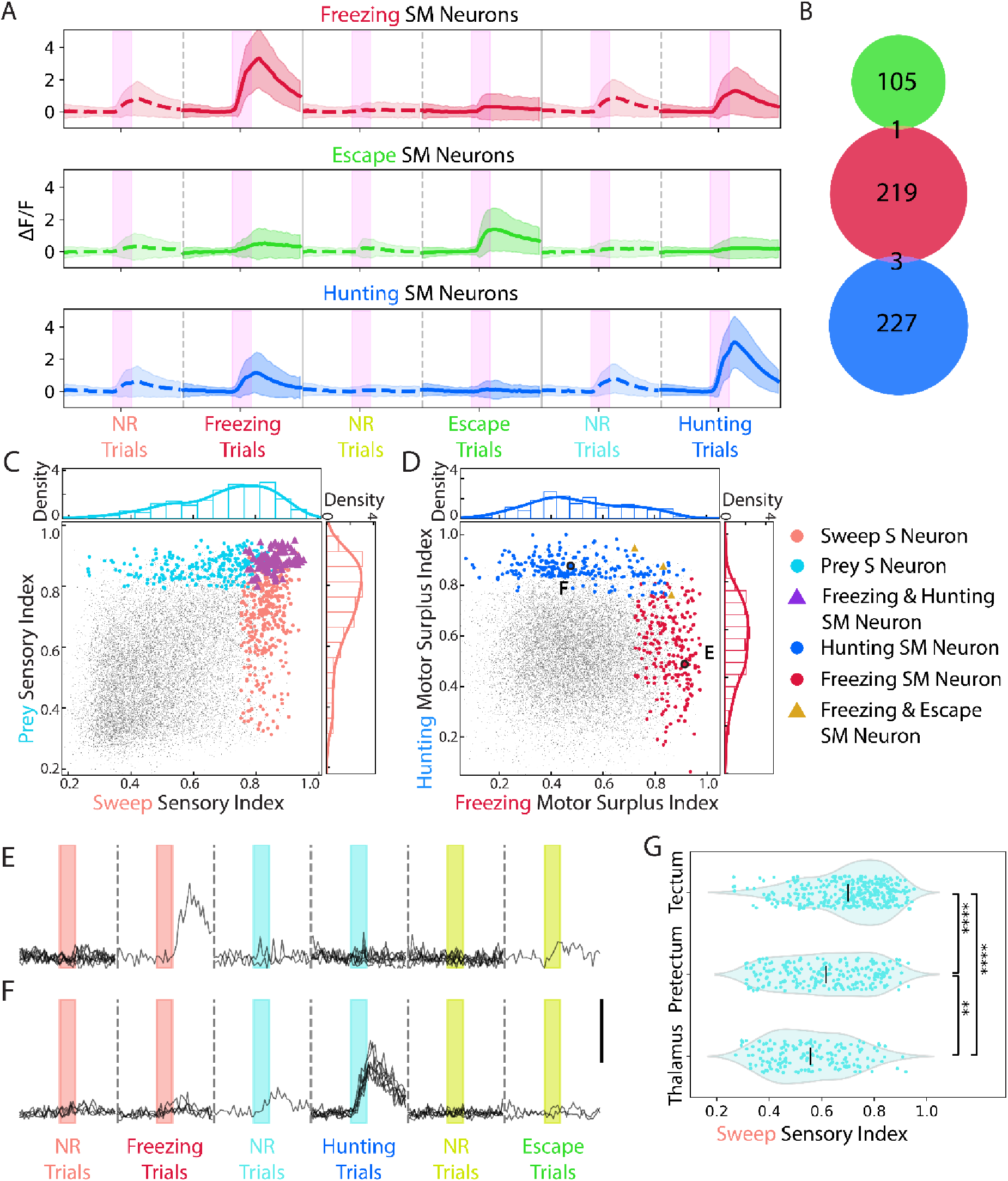
Functional segregation of the different types of sensorimotor neurons in the tectum. (A) Activities of tectal SM neurons during motor response or no response trials. Pink bars represent 4-second stimulus presentation. n = 7 larvae. Shading represents Standard Deviation. (B) Venn diagram of the three SM neuron populations in the tectum. Green = escape SM, red = freezing SM, and blue = hunting SM neurons. (C) The distribution of the tectal sweep (light red) and prey (light blue) sensory neurons on the sweep and prey sensory index. Purple triangles: neurons belonging to both populations. (D) The distribution of tectal freezing (dark red) and hunting SM neurons (dark blue) on the MSI of freezing and hunting. Gold triangle: neuron belonging to both populations. (E-F) Responses of example tectal freezing and hunting SM neurons from D. Scale bar indicates ΔF/F = 3. (G) Distribution of the sweep sensory index of prey sensory neurons in the tectum, pretectum, and thalamus. Black lines indicate averages **, p<0.01. ****, p<0.0001, Kruskal-Wallis test, followed by Dunn’s test.

To look more closely at how the tectum generates separate hunting and freezing sensorimotor populations, we first plotted the functional distributions of the prey and sweep sensory neurons in the tectum, and saw that these populations were highly overlapping, with many prey sensory neurons having a high SI for sweep, and vice versa (figure 6C). In contrast, SM neurons for hunting and freezing had highly divergent responses; freezing SM neurons generally had low MSI for hunting (figure 6D, red), indicating little response during hunting trials (example neuron, figure 6E). Conversely, hunting SM neurons had low MSI for freezing (figure 6D, blue) and were unresponsive during freezing trials (example neuron, figure 6F). How is the tectum able to go from overlapping sensory to divergent sensorimotor populations? In the case of the prey stimulus, other areas such as the pretectum and thalamus also receive input from prey-selective retinal ganglion cells^15^, and contain prey sensory neurons (figure 2C). We plotted the sweep SI for prey sensory neurons in the pretectum and thalamus, found that these neurons are less responsive to the sweep stimulus (figure 6G), suggesting they may be more selective for prey than the prey sensory neurons of the tectum. Thus, these other areas could detect the presence of prey, and provide input to the tectum, which would then combine this input with its precise object location information to drive directed hunting behavior.

Unlike the sweep and prey stimuli, sweep and looming stimuli occupied somewhat different regions of visual space (Supplementary figure 2A), and this could cause less overlap among the sweep and looming sensory neurons in the tectum. However, we did observe neurons that belonged to both sweep and looming sensory populations (Supplementary figure 16A). As was the case for freezing vs. hunting, the escape vs. freezing sensorimotor populations were more diverged than their sensory pairs, despite the fact that both behaviors are responses to predators (Supplementary figure 16B-D). These results suggest that the hunting, freezing, and escape sensorimotor pathways have already diverged at the level of the tectum.

### Downstream areas contain sensorimotor neurons for the three behaviors

To investigate the flow of visuomotor information from the tectum to other areas, we next looked at the anatomical distribution of SM neurons within three areas likely to be downstream of the tectum^29^; the NI, thalamus, and pallium. Within the NI, SM neurons were distributed in distinct patterns, with escape SM neurons localized to the posterior isthmi (figure 7A and Supplementary figure 17A), in a region corresponding to the cholinergic domain (Supplementary figure 17B), suggesting some potential functional specialization for escape behavior. We next looked at the functional properties of the NI SM neurons, by plotting each neuron’s MSI. We found that the escape and freezing SM neurons in this area were generally distinct populations, although there was a small subset of neurons that were classified as both freezing and escape SM neurons (10.7% of all NI freezing SM neurons). There was a smaller population of NI neurons that were both freezing and hunting SM neurons (figure 7C).

**Figure 7:**
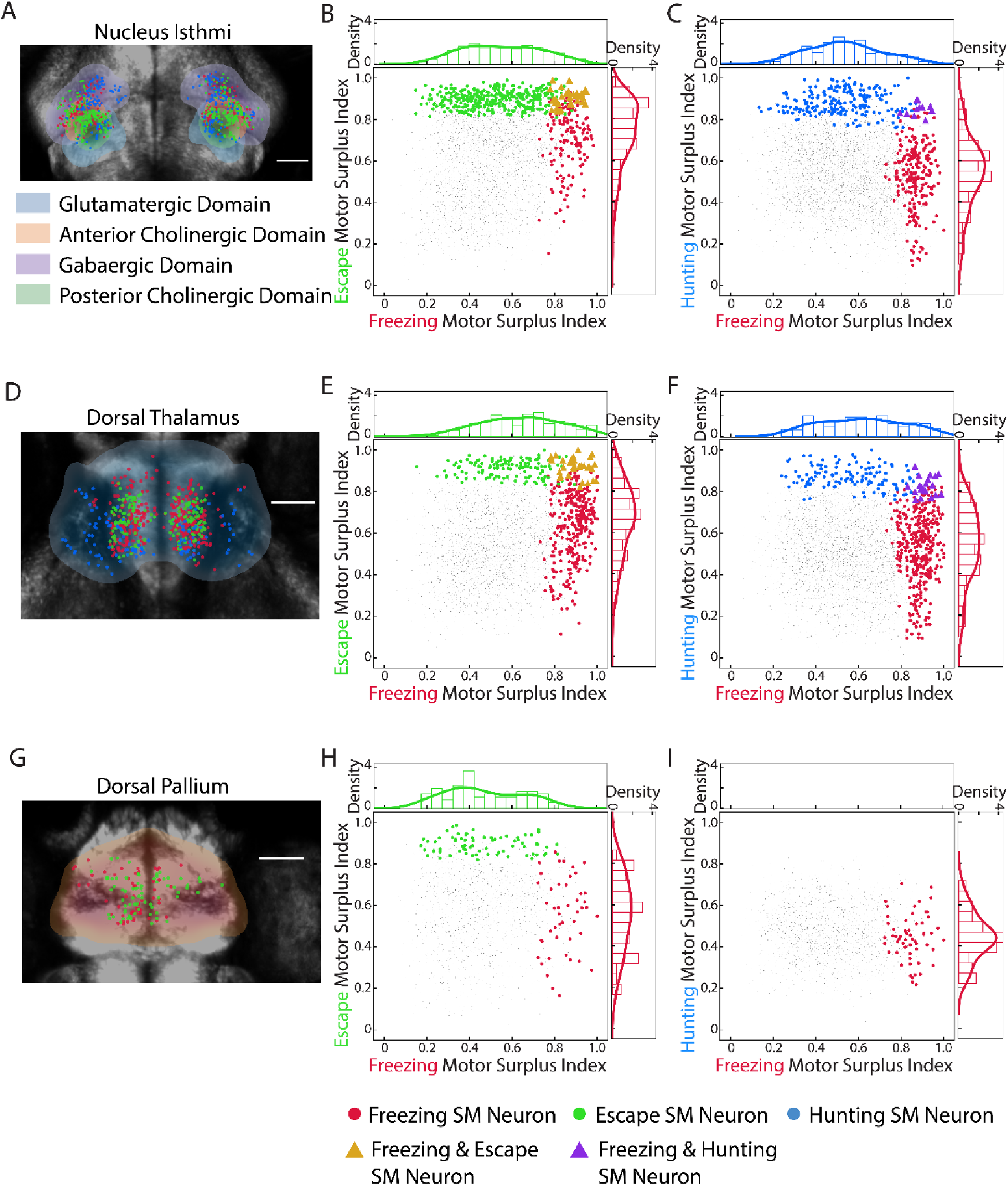
Locations and functional properties of SM neurons in NI, thalamus, and pallium. (A) Dorsal view of the locations of freezing (red), escape (green), and prey (blue) SM neurons in the NI. Shaded regions indicate locations of different domains in the mapZebrain atlas. Scale bar indicates 50µm. (B) The distribution of the NI freezing (red) and escape (green) SM neurons on the MSI of freezing and escape. Golden triangles: neurons belonging to both populations. (C) The distribution of NI freezing (red) and hunting (blue) SM neurons on the MSI of freezing and hunting. Purple triangles: neurons belonging to both populations. (D) Dorsal view of the locations of the SM neurons in the thalamus. (E) Distribution of thalamic freezing and escape SM neurons. (F) Distribution of thalamic freezing and hunting SM neurons. (G) Dorsal view of the locations of the SM neurons in the pallium. (H) Distribution of pallial freezing and ecape SM neurons. (I) Distribution of pallial freezing SM neurons (no hunting SM neurons were identified in the pallium).

The number of SM neurons that are classified as belonging to two pathways will depend on the threshold we use to identify SM neurons. We therefore varied the threshold from 3 to 10%, and found that even with a more lenient threshold, the proportion of neurons classified in multiple pathways remained low (Supplementary figure 18A-C), with the number of freezing + escape neurons going from 5% of the total freezing population at our standard threshold to 13% using the most lenient threshold.

Like the NI, the thalamus also contains all three types of SM neurons, but the freezing and escape populations are restricted to medial thalamus, suggesting this could be a predator-specific region (figure 7D). As in the NI, most SM neurons were specific for one pathway, although there were small populations of freezing + escape and freezing + hunting SM neurons (figure 7E-F).

In contrast to the NI and thalamus, within the dorsomedial pallium, which is the homologue of the mammalian amygdala^52^, we identified only freezing and escape SM neurons (figure 7G). The two populations were entirely separate, with no freezing + escape neurons (figure 7H). Overall, our analysis of these three areas suggests that after the decision to execute one of the three behaviors and subsequent segregation of the pathways in the tectum, the freezing, escape, and hunting pathways continue to be segregated, although there are a few neurons in NI and thalamus that are part of two pathways. The exact functional roles of these areas and dual pathway neurons remain to be determined.

What roles could the SM neurons in different regions be playing in visual behaviors? To begin to explore this question, we asked whether the SM neurons in some regions are more sensory-driven or more behavior correlated than others. We first looked at the distribution of SM neuron SI’s within each area as an indication of how much they were driven by the visual stimulus. The freezing SM neurons in the five main areas all had similarly low SI’s of ∼0.5 for sweep stimuli, although SM neurons in the pallium tended to have a lower SI of ∼0.4 (Supplementary figure 19A). Escape SM neurons also had similar SI’s for looming across all five areas, although pallium and NI indices were slightly lower (Supplementary figure 19B).

Hunting SM neurons across areas also had similar SI’s for prey, although the NI neurons’ indices were somewhat lower (Supplementary figure 19C). We lastly looked at the MSIs of SM neurons in each area as an indication of how correlated their motor surplus was with the visual behavior. One possibility is that downstream areas could be more motor correlated than the tectum, but we found that the MSI for freezing of all the freezing SM neurons was generally similar across all areas, although the thalamus had a slightly higher MSI (Supplementary figure 19D), and the same was true for thalamic escape SM neurons (Supplementary figure 19E). For hunting, the MSI distributions were also similar, although the NI SM neurons had a slightly higher MSI (Supplementary figure 19F). This suggests that the SM neurons we identified for each behavior are generally similar to each other, in terms of having little sensory drive and being equally correlated with behavior, despite being located in five different brain regions.

Overall, the analysis of our functional imaging dataset allowed us to identify sensory neurons for sweep, prey, and looming visual stimuli, which were located primarily in the tectum (Supplementary figure 20A, light red, blue and green dots), and these neurons were broadly tuned. By calculating the MSI for the three behaviors for each neuron in the dataset, we found populations of SM neurons that were correlated with each behavior. These neurons were found in the tectum, pallium, thalamus, and NI (Supplementary figure 20A, dark red, blue, and green dots), and were highly specific for their primary behavior, with little response to sensory stimuli or other visual behaviors.

## Discussion

Here, we developed an imaging-compatible behavioral assay for the innate, visually evoked freezing response in larval zebrafish, and used this assay as well as the established hunting and escape behaviors to map the neural circuits for detection of visual stimuli and selection of visual behaviors. By identifying the sensory and behavior-correlated neurons for hunting, freezing and escape, we can observe how visual sensory responses are transformed into behavioral outputs in the optic tectum, and describe for the first time the functional organization of the downstream areas that receive this input (Supplementary Figure 20).

Not surprisingly, the tectum contained large populations of sensory neurons for all three visual stimuli, and these neurons are likely integrating tectal retinal ganglion cell inputs to respond to moving objects in the visual field. The sensory neurons may then provide input to tectal SM neurons of the corresponding behavior, which could also receive other inputs that reflect fear, satiety, locomotor or other internal states^24,25,53^ to influence the visuomotor decision. One surprising finding was that the SM neurons overall had very little response to the visual stimuli, and minimal overlap with the corresponding sensory population. This is different from the pathway mediating left/right turns^45^, but it may be that freezing, escape and hunting behaviors are very consequential for the animal, and thus the decision to behave is more of an all or nothing one. SM neurons for these three behaviors thus seem to act as AND gates, only firing and perhaps triggering the behavior when both the visual stimulus and the appropriate state information are present.

Another unexpected result was the degree to which the tectal SM neurons were specific for one behavior. One might expect there to be some “predator” neurons, which would respond in both freezing and escape trials. However, the freezing/escape segregation was already apparent in the tectum, suggesting that, at least for these important innate behaviors, the animal is directly deciding whether to freeze or escape, rather than first identifying the stimulus as a predator.

Our findings that the tectum alone in our imaging volume contained both sensory and SM neurons for the three stimuli/behaviors, and that the SM neurons there were already highly correlated with behavior, suggest that the tectum could be the key site of visuomotor decisions on whether to behave. Our partial correlation analysis showed that freezing and hunting SM neurons in the tectum have significant correlations with the corresponding tectal sensory neuron populations, suggesting that these SM neurons are integrating sensory neuron input, perhaps using the tectum’s map of visual space^26^ to determine the location of the object, in order to respond appropriately. However, one caveat is that we cannot infer direct monosynaptic connectivity from this analysis. Interestingly, we do not find the same evidence of within-tectum sensory correlations for escape SM neurons (Figure 5J). It may be that escape is different from the other two behaviors, in that when an animal is hunting an object or freezing in order to evaluate it, knowledge of its position in the visual field is crucial, whereas during escape, the precise predator location may not be as important as executing an evasive movement roughly away from the threat.

The tectum may not be the only area where SM neurons receive sensory neuron input; in sampling the other visual areas with both types of neurons, we found that pretectal hunting SM neurons are significantly correlated with the pretectal prey sensory population (Supplementary figure 15A). Pretectal prey sensory neurons are more specific for prey stimuli than their tectal counterparts (Figure 6G), so the pretectal hunting SM neurons may be using the information from the prey-selective retinal arborization field 7^15^ to identify prey and initiate eye convergence. Pretectal SM neurons could then activate tectal hunting SM neurons, which would integrate the information on object location from the tectal sensory neurons to precisely target prey. Indeed, hunting command-like neurons have been identified in the pretectum, and they have projections to the anterior optic tectum^32^. It is likely that these neurons are part of the pretectal hunting SM population, and we indeed see that a large percentage of pretectal hunting SM neurons are correlated with the tectal SM population (Supplementary figure 15B). These data support a model where identification of prey occurs in the pretectum, and this information is transmitted to tectal hunting SM neurons, which encode precise prey-directed swims based on object location.

In addition to the tectum, we also found significant populations of hunting, freezing and escape SM neurons in the thalamus and NI (homologue of the parabigeminal nucleus), while the dorsomedial pallium (homologue of the amygdala) contained only freezing and escape SM neurons. In terms of escape, our results agree with recent studies in zebrafish that found looming stimuli evoke responses in the tectum, thalamus, and pallium^45,61^, although escape behavior was not recorded in these studies, so escape SM neurons could not be identified. Our findings are also highly consistent with studies in mice where optogenetic stimulation of the thalamus, parabigeminal nucleus and amygdala evoked freezing and escape^33,62^, validating our approach and suggesting that the neural circuits for these important innate behaviors are deeply conserved. In mice, the LP region of the thalamus has been considered to mediate freezing and the parabigeminal nucleus to mediate escape, based on the fact that optogenetic activation of these areas biases animals toward one of the behaviors^33^. In contrast, our data show that these areas contain a significant number of SM neurons for both defensive behaviors (pallium) or all three visual behaviors (thalamus and NI). It may be that, rather than mediating a single behavior as the optogenetics results might suggest, these areas are involved in functions relevant to multiple behaviors, such as stimulus selection in the case of the NI^57^, or object identification in the case of the thalamus^30^. However, one caveat is that we do not know the effect of the SM neurons we identified, and some subset of them may be inhibitory. Thus, the exact role of these areas in these behaviors remains to be determined.

In addition to sensory and SM neurons, one further component of the complete stimulus to action pathway would be motor neurons. These neurons may be downstream of the SM neurons, and should respond during behaviors regardless of whether they are evoked by visual stimuli or some other pathway. For example, the escape SM neurons in the tectum could provide input to the prepontine neurons that have been shown to mediate auditory escape^64^. We did not evoke the three behaviors with other sensory modalities, and our dataset does not include enough “spontaneous” or non-stimulus-evoked behavior to distinguish motor neurons from SM neurons, but in the future it will be interesting to identify this third type and its relationship with SM neurons.

In summary, we found that the innate visual freezing response, as defined by immobility and bradycardia, can be triggered by a sweeping visual stimulus in zebrafish larvae. We identify sensory neurons for prey, sweeping and looming stimuli primarily in the tectum, and hunting-, freezing-, and escape-correlated neurons in the tectum and downstream areas. Within the tectum, freezing and hunting SM neurons are correlated with their corresponding sensory neurons populations, suggesting that they combine visual and behavioral state information to produce a behavioral output. The SM neurons in the thalamus, nucleus isthmi and dorsomedial palium are likely activated by SM output neurons of the tectum, and may play roles in the identification and selection of prey or predatory stimuli that evoke hunting, freezing and escape.

## Acknowledgements

Funding was provided by the Hong Kong Research Grant Council (16103522, 16101221, 16103118, and 26100617) and the Chau Hoi Shuen Foundation. We would like to thank Yukinori Hirano and members of the Semmelhack lab for feedback on the manuscript.

## Author Contributions

Conceptualization, P.Z., Y.T. and J.L.S. Methodology, Y.T. Software, P.Z., Y.T. and T.K.C.L. Formal Analysis, P.Z., Y.T. and Y.H. Investigation, P.Z., Y.T., G.T., and K.K.Y.C. Resources, I.P.L. and B.K. Writing – Original Draft, J.L.S. Writing – Review & Editing, all authors. Supervision, J.L.S. Funding Acquisition, J.L.S.

## Supplementary Video Legends

**Supplementary Video 1: Sweep stimulus suppresses spontaneous swims.** Swimming was recorded from above at 200 frames per second. The stimulus was a 15° diameter dark disk moving at 60°/second on a red background.

**Supplementary Video 2: Sweep stimulus causes a reduction in heart rate.** The stimulus was a 15° diameter dark disk moving at 60°/second on a red background. Heart rate was recorded from the side at 100 frames per second.

**Supplementary Video 3: Prey stimulus triggers hunting behavior.** Eye and tail movements recorded from above at 200 frames per second, in response to a 4° diameter UV dot moving at 120°/second for 6 seconds. For imaging experiments, the speed was 60°/second and duration was 4 seconds.

**Supplementary Video 4: Looming stimulus triggers escape behavior.** Tail movements were recorded from above at 200 frames per second, in response to a dark disk expanding to 60° in diameter.

## Code Availability

Key code, including neuronal data preprocessing, SI and MSI calculation, sensory and sensory motor neuron selection, and behavioral analysis pipeline (heart rate analysis, bradycardia detection, hunting detection, tail bout detection, and escape classification) is available on github: https://github.com/SemmelhackLab/Freezing_Code/tree/main

## Methods

### Animals

Zebrafish larvae were raised in Danieau’s solution in a petri dish inside an incubator set to a standard 14:10 hour light cycle at 28.5°C. All the larvae used in this study carried homozygous *mitfa* skin-pigmentation mutation (nacre). Tg(elavl3:H2B-GCaMP6s) larvae were used for the imaging experiments. All animal experiments were conducted with approval from the Animal Ethics Committee of Hong Kong University of Science and Technology.

### Behavioral Experiments

Each larva was embedded at 5 or 6 days post fertilization in 2% low melt agarose on a Rinzl plastic coverslip (Delta Microscopies). Larvae that were embedded on day 6 were fed 1 mL of a dense paramecium culture per petri dish on day 5. After solidification, the agarose around the head and tail was then removed to free the eyes and the caudal half of the tail. The embedded larvae were allowed to accommodate overnight before the experiment.

Behavioral experiments were conducted in a chamber where stimuli were projected on one wall and high-speed cameras recorded from above and below, as previously described^39,40^. Before the experiment, the coverslip with the larva was inserted into a 3D printed resin scaffold connected to an XYZ manipulation stage (Heng Yang Optics). The larva was then secured inside an arena filled with Danieau’s solution. The arena is a rectangular tank made of transparent acrylic board, with the frontal wall made of Teflon fabric serving as a screen. The dimensions of the arena were 60×80×30 mm. The distance between the screen and the fish was 10 mm. The top camera (Photon Focus, MV1-D1312E-160-CL-12) recorded the eye and tail movement of the fish, while the left camera (HIK Robot MV-CA013-21UM/UC 130) captured the heart rate. The frame rate was 200 frames per second for the top camera and 100 frames per second for the bottom camera. Recording from the two cameras and the stimulus display were synchronized with custom Python scripts. For both the behavioral and imaging experiments, a customized LightCrafter (DLP® Light Crafter™ 4500 modified by EKB Technologies Ltd was used for stimulus projection, placed 13 cm away from the frontal screen. The wavelengths of the LEDs were 385, 415, 470, 590 nm for UV, blue, green, and red, respectively. The light intensity of the stimulus or the colored background was 886 nW/cm^2^, consistent across colors.

Before the start of stimulus presentation, the fish were allowed to accommodate for 5 minutes. Stimuli were presented in pseudo-random order. After each stimulus display, there was a trial with no stimulus, and the inter-trial interval was 1 minute. The duration of each trial was 22 seconds, and the stimulus was shown during the 9^th^ -13^th^ second, unless specified otherwise.

### Visual Stimuli

Visual Stimuli were designed in Python using PsychoPy^37^. For the behavioral experiments to optimize the sweep stimulus (figures 1 and 2), the stimulus was a dark disk moving horizontally back and forth across the red background (-60° to 60° in azimuth). The starting azimuth of the stimulus was either -60° or 60° alternating among trials to avoid habituation. The elevation of the stimuli was 5° above the midline. The diameter of the stimulus was 15°, and the speed was 60° per second, unless otherwise stated.

For the imaging experiments, the sweep stimulus was a dark disk with a diameter of 15° and a speed 60°/s, on a red background. The prey stimulus was a 4° UV dot with the same horizontal movement trajectory as the sweeping stimulus, but an elevation of 15° to promote hunting behavior. The looming stimulus was a dark expanding disk on the same red background, starting from 6° and ending in 60° in diameter. The expansion size-to-speed ratio (l/v) was 60ms. After reaching the maximum size, the looming stimulus remained stationary until the 4 second stimulus period was over. The looming disk was presented in the center of the visual field (azimuth = 0°) and the elevation of the center of the disk was -20° (Supplementary figure 2A), to promote escape responses. All trials lasted for 22 seconds in total. We presented eight sweep stimuli, followed by eight prey stimuli, followed by eight looming stimuli, with an interstimulus interval of 2 minutes.

### Heart Rate Analysis

To analyze the heart rate, an ROI of around 100×100 pixels that contained the heart was manually cropped, and rhythmic pixels within that ROI were selected (figure 1B, left and middle). The first criterion for rhythmic pixels was having stable local maximum and minimum intensity values (each local max>(.75* last local min + .25*last local max), and each local min<(.75*last local max +.25*last local min)), and the second was having a stable heart rate over the whole trial (standard deviation of less than 0.55). If the number of rhythmic pixels was less than 5%, the trial would be excluded from further analysis. The intensity trace of each pixel was gaussian filtered (figure 1B, right), and the heart rate for that pixel was calculated as the reciprocal of the inter-beat interval. Heart rates from all the rhythmic pixels were averaged to calculate the heart rate for the trial. Heart rate was normalized by dividing by average heart rate in the 6 seconds before stimulus onset.

### Bradycardia Identification

To identify bradycardia, a dynamic ceiling, or local presumptive max heart rate, was calculated^38^. The ceiling was acquired by applying a maximum filter with a 300-frame window to the median filtered heart rate trace. The distance between the original heart rate and the ceiling, termed heart-rate-to-ceiling, was calculated for every frame to examine the local decrease in heart rate. If the heart-rate-to-ceiling of the frame was larger than 3 standard deviations of the heart-rate-to-ceiling of all the frames from no stimulus trials, the frame was considered a bradycardia frame. When the bradycardia persisted for 50 frames (250 ms), the period was annotated as a bradycardia episode.

### Eye and Tail Tracking

Eye and tail tracking were conducted by Ztrack in Python. In brief, the contours of both eyes were found by multi-threshold binarization. The binocular eye angle was calculated from the image moment of the contours. To detect eye convergence, the distribution of binocular eye angles was fitted with a kernel density estimation and the location of the first local maximum within 20-50° was chosen as the threshold for eye convergence episodes of the corresponding fish. If no local minimum was found, trials were checked manually for convergence.

For tail tracking, a skeleton of tail with 20 nodes was tracked from the binarized image. The tail curvature was determined as the average of the nodal angle of the last 10 nodes. To detect swimming bouts, hysteresis thresholding was applied to the derivative of the tail curvature with respect to time. Bouts near each other would be merged as one bout and bouts that were less than 8 frames were discarded. The swimming probability of each frame was the probability that a swimming bout was detected at that point across all trials.

### Escape Classification

To identify escape bouts, all swimming bouts detected in the dataset were processed by Principal Component Analysis (PCA), followed by K-Means Clustering. 8 features of each swimming bout were dimensionally reduced by PCA: the curvature, angle of the tail tip point, angle of the tail middle point, mean and max velocity of tail movement, frequency of tail movement, duration of the bout, and the integral of the tail bouts. The first 3 PCs explained over 90% of the variance. K-means clustering was applied to the first 3 PCs of all tail bouts and 5 clusters were formed. The cluster with high frequency, high velocity, and short bout duration was selected as escape bout cluster. To examine the accuracy, 80 new bouts were manually annotated, and the classification showed 92% accuracy compared to manual annotation.

### Behavioral Categorization

The behavior of fish in each trial was classified into one of 5 types: Freezing, Hunting, Escape, Spontaneous Swim, and No Response (figure S3B). To be classified as one type of behavioral response, phenotypical signatures of the behavior had to be detected, and the behavior had to start during the 4 seconds after stimulus onset. The classification criteria are shown in Supplementary figure 3A. To be classified as a freezing trial, there must be a bradycardia episode (heart-rate-to-ceiling larger than 3 SD for 250 ms, Supplementary figure 3A, upper panel) starting during the stimulus period, and no eye or tail movement. For escape, an escape bout (high frequency and velocity, within the escape cluster, Supplementary figure 3A, middle panel) starting during the stimulation period was required. For hunting, eye convergence above the first local minimum of eye angle for the animal (Supplementary figure 3A, lower panel) starting during the stimulus period was required. For spontaneous swim, there had to be a non-escape bout with no escape bout or eye convergence during the stimulation. Finally, trials with none of the above types were considered no response trials. Trials with a mixture of behaviors (e.g. freezing followed by escape) during the stimulus period were excluded in further analysis.

### Calcium Imaging Acquisition

2-photon calcium imaging was conducted with a Nikon upright 2-Photon microscope (Nikon A1 MP multiphoton) with a custom-built behavior setup. To achieve multi-plane imaging, an Electrically Tunable Lens (Optotune) was added between the microscope and the 20x Nikon Objective. The ETL was synchronized with the imaging system and triggered every frame. The wavelength of the laser was set to 920 nm. Images of 512 x 512 pixels were recorded at 2 volumes/second from 14 planes, with around 10 microns between planes.

The imaging chamber was similar to that used in the behavioral experiments. The fish, embedded on plastic cover slip and secured to the XYZ stage, was placed 1.5 mm beneath the objective front lens. Both the front lens and the fish were immersed under water held in a rectangular arena made of transparent acrylic board, except the front wall made of Teflon fabric and serving as the screen. The distance between the screen and the fish was 10 mm, and the screen area was 70° high x 140° wide, with the larva positioned at 0°. A LightCrafter (DLP Lightcrafter 4500 MKII) was placed in front of the screen, 15 cm away, to project the virtual stimulus. Two sets of cameras were placed around the arena. A bottom-view camera (Photonfocus, MV1-D1312E-160-CL-12) and light record the eyes and tail at 200 frames per second. A side camera (HIK Robotics, MV-CA013-21UM) recorded the heart rate of the fish at 100 frames per second.

The presentation of the stimulus, the recording of cameras, and the tuning of the ETL were all synchronized with the calcium imaging session by Labjack (Labjack U3), custom-built circuits, and custom Python scripts.

### Calcium Imaging Analysis

All calcium imaging analyses were performed with custom-written Python code. Motion correction and neuron segmentation were conducted with suite2p^43^. Volumetric imaging data from three fish was used to train a suite2p classifier to segment ROIs. For each plane, around 800-3000 ROIs were segmented, and ROIs larger than 300 *μm*^2^ or smaller than 30 *μm*^2^ were discarded. ΔF/F of each neuron were calculated as (F(t)-F_0_)/F_0_ in each trial, where F_0_ was calculated as the average intensity over the 4 seconds before stimulus onset, and F(t) was the intensity trace of each neuron.

All calcium imaging registration was done with Advanced Normalization Tools (ANTs). Software and all reference volumes were downloaded from mapZebrain^44^. The registration was performed in 3 steps: 1) 3d registration from the z-stack to the atlas. 2) 2d registration from the calcium imaging video to the z-stack. The average calcium image was used as the moving image and the reference image was the best-matched plane chosen from the z-stack by template matching (OpenCV). 3) Transformations were applied to the coordinates of segmented neurons. Anatomical regions of each registered neuron were identified by the masks downloaded from mapZebrain.

### Selection of Active Neurons

Neurons with either sensory- or motor-related activity were selected from all the segmented cell bodies using linear regression. Sensory-relevant activity was represented by a *sensory regressors* (Regressor_S_ , one for each stimulus) built by a box car function over the stimulus presentation window (9-13s), then convolved with the GCaMP6s kernel (i.e., an exponential decay 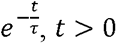 with a time constant *τ* = 7s^65^). The motor-related activity was similarly represented by *visuomotor regressors,* Regressor_SM_ . To construct a visuomotor regressor, we build a 1 second box car function starting from the behavior onset (i.e., bradycardia without movement, escape tail bouts, and eye convergence movements as the onsets of freezing events, escape events and hunting events, respectively). We then shifted the box car by -0.5 seconds and convolved it with the GCaMP6s kernel^65^. Only motor events that started within the stimulus presentation window (9-13s) were used to construct the visuomotor regressor. For each neuron, we regress its activity in each trial to one of the sensory and motor regressors. Neurons with at least 30% of trials having a slope>0.35 and R^2^ >0.36 to any sensory or visuomotor regressor were considered active neurons. In the end, we found 34% of segmented cell bodies were classified as active neurons and used this set for subsequent analysis.

### Sensory Index Calculation and Sensory Neuron Identification

We used a similar method to identify sensory neurons as in Chen *et. al*.^45^ In brief, if the information encoded in a neuron is purely sensory, its response to the repeatedly presented stimulus across trials should be similar and thus having a highly periodic activity. Therefore, we can select a sensory neuron using periodicity as a sensory index, which measures how similar the neuron’s responses are across trials.

To identify sensory neurons for a given stimulus, we first selected neurons that responded to the stimulus within the population of active neurons, using the same Regressor_S_, and took the top 10% most correlated neurons (Pearson’s correlation coefficient). To calculate the periodicity, we form a Trace_avg_ by tiling or concatenating a neuron’s trial-average response to the stimulus over each of the trials (figure 2B). Then the periodicity, i.e., the sensory index (SI), was calculated as the square root of the variance ratio between Trace_avg_ and the original activity Trace (see equations below). The periodicity of each neuron was calculated per stimulus; thus, each neuron had 3 sensory indices.

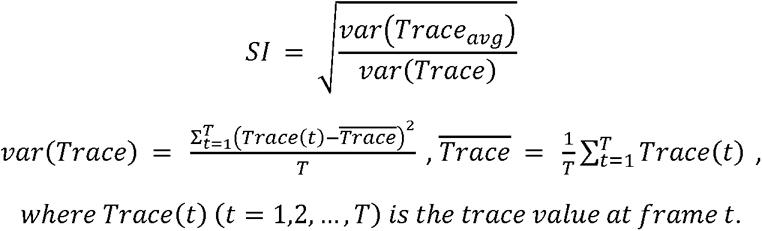

We define the top 15% of neurons by sensory index as sensory neurons. The selection was conducted for each stimulus, and one neuron could be selected as a sensory neuron for more than one stimulus. Neurons that were not spatially colocalized across fish were eliminated by a *Spatial colocalization test,* similar to Spatial p-value filtering^46^. Briefly, for a given neuron *i,* we compute the average distance of this neuron to the nearest neuron (other than itself) within the same category in each fish. This average distance is then compared with the null distance. The null distance is the distance of neuron *i* to a neuron randomly sampled from all the active neurons combined across all fish (recall that the fish is registered to a standard brain atlas). The random sampling of neuron and the null distance were repeated 5000 times to build a null distribution, which is then fitted to a normal distribution. The neuron *i* is considered anatomically colocalized across fish if the average nearest distance falls under the 2.5% normal distribution percentile (i.e. p-value < 0.025).

### Motor Surplus Index Calculation and Sensorimotor Neuron Identification

The calculation of Motor Surplus Index was inspired by the motor decomposition method in Chen et al.^45^. Firstly, for each active neuron, a trace surplus was calculated by subtracting the average trace from the original trace (figure 3C). Then, the Pearson’s correlation coefficient between the trace surplus and the visuomotor regressor (Regressor_SM_ , same as in “**Selection of Active Neurons**” but concatenated across trials) was calculated. The correlations are linearly scaled to a motor surplus index (MSI) between 0 and 1, using the min and max correlation in each fish. Note that this scaling is only for better visualizing results across fish, but has no effect on sensorimotor neuron identification. The MSI of each neuron was calculated per behavioral response (freezing, escape, and hunting) using recordings during the corresponding stimulus window (sweep, looming, and prey), resulting in 3 MSIs for each neuron. For each behavioral response and each fish, the top 3% of the neurons with the highest MSI are selected as the preliminary SM neurons. The Spatial colocalization test (as described in “**Sensory Index Calculation and Sensory Neuron Identification**”) was applied to further select neurons that are anatomically colocalized across fish (i.e., with a p-value < 0.025) and were used as the final SM neurons for further analysis. Out of all the preliminary SM neurons for each behavior, 74.5%, 74.3% and 74.3% of them passed the Spatial colocalization test.

### Behavior Prediction using Freezing SM Neurons

The behavior prediction using freezing SM neurons was conducted on a leave-one-out basis. We held out one trial and used the remaining trials to select SM neurons based on the top 3% MSI criteria. The behavior prediction is determined by comparing the activity of selected SM neurons in the held-out trial with the average activity in behavioral trials and in non-behavioral trials. In particular, we use the maximum of the average activity trace in a trial as the feature. If this maximum in the held-out trial is closer to the maximum value of the behavioral-trial average, we will predict the held-out as a behaviorial trial, and vice versa. Under this method, the held-out prediction could only be conducted when there are at least one behavioral and one non-behavioral trials after held-out. Therefore our results are from a subset of held-outs for certain fish. For example, for a total of 8 sweep trials in one fish, if there was only one behavioral trial, then there would be 7 held-outs that we can make the prediction.

The prediction accuracy for each possible held-out trial (100% or 0%, left column, Supplementary figure 9D and 10) is compared per held-out trial with the accuracy using *null SM neurons.* Null SM neurons were selected similarly, using sensorimotor regressors built according to trial-permuted behavior labels, for the activity in the seven remaining trials. Since there are multiple ways to permute the behavior labels, a different set of null SM neurons will be selected for each permutation. For example, if there were 8 trials in total and 2 trials are behavioral trials, if the held-out trial is a behavioral trial, there would be 7 behavior label permutations and sets of null SM neurons. If the held-out trial is a non-behavioral trial, there would be 21 permutations.

For each permutation and corresponding set of null SM neurons, we use the same method of trace maximum comparison as for the real SM neurons to make the behavior prediction for the held-out trial. We then average this accuracy across permutations to get the null prediction accuracy (right column, Supplementary figure 9D and 10).

### Single-photon optogenetic stimulation

Zebrafish larvae were embedded with eye and tail were freed, and placed in the behavioral chamber. An optical fiber, 50 µm (M14L01 Thorlabs) or 105 µm in diameter (M15L01 Thorlabs), was held about 0.5mm above the head and guided the light from a blue light fiber-coupled LED (M470F4, Thorlabs). We used 0.06 mW or 0.26 mW light power, measured at the fiber tip. The position of the optic fiber was controlled using an XYZ manipulation stage (Heng Yang Optics). We targeted the anterior and posterior tectum and stimulated each spot with 10 trials. Each trial contained a 5 s stimulation period and a 5 s interval between each trial. Behavior was recorded at 200 fps and manually annotated for eye convergence and escape tail movements.

### Partial Correlation Calculation between Behavior Residuals of Sensory and Sensorimotor Neurons

In each fish, we used response during the presentation of a given stimulus concatenated across trials to represent the activity of each sensory neuron or SM neuron. For example, if there are 8 sweeping stimulus trials and 8 frames during each stimulus presentation. *S* is a [64 × *m*] matrix and *SM* is a [64 × *m*] matrix, and *n* and *m* are the numbers of sensory and SM neurons, respectively. Neurons that are categorized as both S and SM are excluded from this analysis (between 0-16.7%, or 1-3 neurons across fish) to avoid issues due to self-correlated pairs in later analysis.

For the concatenated response of a sensory neuron *S_i_* (i=1,2,…,n) and a SM neuron *SM_j_* (j=1,2,…,m), i.e., a column in the above S or SM matrices, the behavior average and non-behavior average are calculated by averaging the responses of behavioral and no-behavioral trials, respectively (Supplementary figure 12A). *S_i,res_* and *SM_j,res_* was calculated by subtracting the behavior average and non-behavior average for each trial, according to its behavior type, from the original responses *S_i_* and *SM_j_* (Supplementary figure 12A). For a pair of S neuron *i* and SM neuron *j*, the partial correlation *ρ*(*S_i_*, *SM_j_*|*B*) is the Pearson’s correlation coefficient between *S_i,res_* and *SM_j,res_*^51^.

### Testing Significant Partial Correlations between Sensory and Sensorimotor Neurons Given Behavior

For each SM neuron *j* in optic tectum (or nucleus isthmi, NI), we quantified its correlation strength with tectal sensory neurons by the 90^th^ percentile of the partial correlations *ρ*(*S_i_*, *SM_j_*|*B*) (Supplementary figure 12C, right). To obtain a null distribution of these partial correlations, we permute the *SM_j,res_* trace across trials, while keeping the 8-frame temporal structure within each trial (Supplementary figure 12B). There are thus in total *k!* possible permutations, where *k* is the number of trials. After each permutation, we similarly compute the partial correlations between the permuted *SM_j,res_* and all the tectal sensory neurons, and calculate a 90^th^ percentile (Supplementary figure 12C, left). Collecting such partial correlation percentiles across the *k*! permutations gives a null distribution for the SM neuron’s correlation strength (Supplementary figure 12D) and an empirical p-value can be computed as *x/k!*, where *x* is the number of permutations resulting in a partial correlation 90^th^ percentile that is equal or larger than to the percentile for the original *SM_j,res_* . The p-values of SM neurons (in the same brain region) from all fish are aggregated and adjusted together for multiplicity into q-values to control the False Discovery Rate (FDR) using the Benjamini-Hochberg procedure. We quantified the percentage of SM neurons with significant q-values (<0.05) in figure 5k. Percentage-by-chance is calculated as a control by using the trial-permuted *SM_res_* in place of the real ones in the above analysis steps. We collect such percentage-by-chance values over 1000 permutations. The 95^th^ percentile of these 1000 percentage-by-chance values is 0 for all three types of SM neurons from both optic tectum (OT) and nucleus isthmi (NI).

To evaluate the potential effect due to shared optical noise, we calculated the percentage of tectal SM neurons with significant partial correlation to a surrogate tectal sensory population (S’ neurons) in Supplementary figure 14. To obtain the S’ neurons, we replace each tectal S neuron with the nearest neuron in the same imaging plane with an SI +/-0.05 from the average SI of tectal neurons in that fish. Average SI for tectal neurons ranged from 0.39-0.58 in the seven fish. The average distance between each S neuron and its S’ surrogate for sweep, looming, and prey stimulus are 6.2 (± 2.7), 11.0 (± 9.1), 7.5 (± 4.9) µm (Mean ± SD), respectively. In Supplementary figure 14B, we selected surrogate pretectal SM’ neurons in the same imaging plane and with an MSI +/-0.05 from the average value for pretectal neurons in that fish. Average hunting MSI was in the range of 0.53 - 0.60 for pretectal neurons in the seven fish.

### Statistics

Figure 1D, S1B, C: Mann-Whitney U test.

Figure 1G, Figure 6G, S1E, G, H, I, S19: The Kruskal-Wallis test, followed by Dunn’s test (Bonferroni correction)

Figure 5J, S9, S10, S11, S13, S14, S15: One sided Wilcoxon signed-rank test.

P-value <= 0.0001 was denoted by “****”

P-value <= 0.001 was denoted by “***”

P-value <= 0.01 was denoted by “**”

P-value <= 0.05 was denoted by “*”

## References

1. Bianco, I.H., Kampff, A.R., and Engert, F. (2011). Prey capture behavior evoked by simple visual stimuli in larval zebrafish. Front. Syst. Neurosci. 5. 10.3389/fnsys.2011.00101.

2. Hoy, J.L., Yavorska, I., Wehr, M., and Niell, C.M. (2016). Vision drives accurate approach behavior during prey capture in laboratory mice. Curr. Biol. CB 26, 3046–3052. 10.1016/j.cub.2016.09.009.

3. Borla, M.A., Palecek, B., Budick, S., and O’Malley, D.M. (2002). Prey capture by larval zebrafish: evidence for fine axial motor control. Brain. Behav. Evol. 60, 207–229. https://doi.org/66699.

4. Eilam, D. (2005). Die hard: A blend of freezing and fleeing as a dynamic defense— implications for the control of defensive behavior. Neurosci. Biobehav. Rev. 29, 1181–1191. 10.1016/J.NEUBIOREV.2005.03.027.

5. Pereira, A.G., and Moita, M.A. (2016). Is there anybody out there? Neural circuits of threat detection in vertebrates. Curr. Opin. Neurobiol. 41, 179–187. 10.1016/J.CONB.2016.09.011.

6. Card, G., and Dickinson, M.H. (2008). Visually Mediated Motor Planning in the Escape Response of Drosophila. Curr. Biol. 18, 1300–1307. 10.1016/J.CUB.2008.07.094.

7. Dunn, T.W., Gebhardt, C., Naumann, E.A., Riegler, C., Ahrens, M.B., Engert, F., and Del Bene, F. (2016). Neural Circuits Underlying Visually Evoked Escapes in Larval Zebrafish. Neuron 89, 613–628. 10.1016/J.NEURON.2015.12.021.

8. Temizer, I., Donovan, J.C., Baier, H., and Semmelhack, J.L. (2015). A Visual Pathway for Looming-Evoked Escape in Larval Zebrafish. Curr. Biol. CB 25, 1823–1834. 10.1016/j.cub.2015.06.002.

9. Yilmaz, M., and Meister, M. (2013). Rapid innate defensive responses of mice to looming visual stimuli. Curr. Biol. CB 23, 2011–2015. 10.1016/j.cub.2013.08.015.

10. Barrios, N., Farias, M., and Moita, M.A. (2021). Threat induces cardiac and metabolic changes that negatively impact survival in flies. Curr. Biol. 10.1016/J.CUB.2021.10.013.

11. Hagenaars, M.A., Oitzl, M., and Roelofs, K. (2014). Updating freeze: Aligning animal and human research. Neurosci. Biobehav. Rev. 47, 165–176. 10.1016/J.NEUBIOREV.2014.07.021.

12. Matsuda, K., Yoshida, M., Kawakami, K., Hibi, M., and Shimizu, T. (2017). Granule cells control recovery from classical conditioned fear responses in the zebrafish cerebellum. Sci. Rep. 2017 71 7, 1–13. 10.1038/s41598-017-10794-0.

13. Mann, K.D., Hoyt, C., Feldman, S., Blunt, L., Raymond, A., and Page-Mccaw, P.S. (2010). Cardiac response to startle stimuli in larval zebrafish: sympathetic and parasympathetic components. Am J Physiol Regul Integr Comp Physiol 298, 1288–1297. 10.1152/ajpregu.00302.2009.-Central.

14. Nordström, K. (2012). Neural specializations for small target detection in insects. Curr. Opin. Neurobiol. 22, 272–278.

15. Semmelhack, J.L., Donovan, J.C., Thiele, T.R., Kuehn, E., Laurell, E., and Baier, H. (2014). A dedicated visual pathway for prey detection in larval zebrafish. eLife 3, e04878. 10.7554/eLife.04878.

16. Simmons, P.J., Rind, F.C., and Santer, R.D. (2010). Escapes with and without preparation: the neuroethology of visual startle in locusts. J. Insect Physiol. 56, 876–883. 10.1016/j.jinsphys.2010.04.015.

17. De Franceschi, G., Vivattanasarn, T., Saleem, A.B., and Solomon, S.G. (2016). Vision Guides Selection of Freeze or Flight Defense Strategies in Mice. Curr. Biol. 26, 2150–2154. 10.1016/J.CUB.2016.06.006.

18. Keleş, M.F., and Frye, M.A. (2017). Object-Detecting Neurons in Drosophila. Curr. Biol. 27, 680–687. 10.1016/J.CUB.2017.01.012.

19. Tanaka, R., and Clark, D.A. (2020). Object-Displacement-Sensitive Visual Neurons Drive Freezing in Drosophila. Curr. Biol. 30, 2532–2550.e8. 10.1016/J.CUB.2020.04.068.

20. Wang, F., Li, E., De, L., Wu, Q., and Zhang, Y. (2021). OFF-transient alpha RGCs mediate looming triggered innate defensive response. Curr. Biol. 31, 2263–2273.e3. 10.1016/j.cub.2021.03.025.

21. Gollisch, T., and Meister, M. (2010). Eye smarter than scientists believed: neural computations in circuits of the retina. Neuron 65, 150–164.

22. Ache, J.M., Namiki, S., Lee, A., Branson, K., and Card, G.M. (2019). State-dependent decoupling of sensory and motor circuits underlies behavioral flexibility in Drosophila. Nat. Neurosci. 22, 1132–1139. 10.1038/s41593-019-0413-4.

23. Zacarias, R., Namiki, S., Card, G.M., Vasconcelos, M.L., and Moita, M.A. (2018). Speed dependent descending control of freezing behavior in Drosophila melanogaster. Nat. Commun. 9, 3697. 10.1038/s41467-018-05875-1.

24. Filosa, A., Barker, A.J., Dal Maschio, M., and Baier, H. (2016). Feeding State Modulates Behavioral Choice and Processing of Prey Stimuli in the Zebrafish Tectum. Neuron 90, 596– 608. 10.1016/j.neuron.2016.03.014.

25. Lenzi, S.C., Cossell, L., Grainger, B., Olesen, S.F., Branco, T., and Margrie, T.W. (2022). Threat history controls flexible escape behavior in mice. Curr. Biol. 32, 2972–2979.e3. 10.1016/j.cub.2022.05.022.

26. Isa, T., Marquez-Legorreta, E., Grillner, S., and Scott, E.K. (2021). The tectum/superior colliculus as the vertebrate solution for spatial sensory integration and action. Curr. Biol. CB 31, R741–R762. 10.1016/j.cub.2021.04.001.

27. Li, C., Kühn, N.K., Alkislar, I., Dublanc, A.S., Zemmouri, F., Paesmans, S., Reinhard, K., and Farrow, K. (2022). Pathway-specific inputs to the superior colliculus support flexible triggering of innate behaviors. Preprint at bioRxiv, 10.1101/2022.07.08.499294 https://doi.org/10.1101/2022.07.08.499294.

28. Hafed, Z.M., Hoffmann, K.-P., Chen, C.-Y., and Bogadhi, A.R. (2023). Visual Functions of the Primate Superior Colliculus. Annu. Rev. Vis. Sci. 9, 361–383. 10.1146/annurev-vision-111022-123817.

29. Wu, Q., and Zhang, Y. (2023). Neural Circuit Mechanisms Involved in Animals’ Detection of and Response to Visual Threats. Neurosci. Bull. 39, 994–1008. 10.1007/s12264-023-01021-0.

30. Branco, T., and Redgrave, P. (2020). The Neural Basis of Escape Behavior in Vertebrates. Annu. Rev. Neurosci. 43, 417–439. 10.1146/annurev-neuro-100219-122527.

31. Fernandes, A.M., Mearns, D.S., Donovan, J.C., Larsch, J., Helmbrecht, T.O., Kölsch, Y., Laurell, E., Kawakami, K., dal Maschio, M., and Baier, H. (2021). Neural circuitry for stimulus selection in the zebrafish visual system. Neuron 109, 805–822.e6. 10.1016/J.NEURON.2020.12.002.

32. Antinucci, P., Folgueira, M., and Bianco, I.H. (2019). Pretectal neurons control hunting behaviour. eLife 8. 10.7554/eLife.48114.

33. Shang, C., Chen, Z., Liu, A., Li, Y., Zhang, J., Qu, B., Yan, F., Zhang, Y., Liu, W., Liu, Z., et al. (2018). Divergent midbrain circuits orchestrate escape and freezing responses to looming stimuli in mice. Nat. Commun. 2018 91 9, 1–17. 10.1038/s41467-018-03580-7.

34. Yu, H., Xiang, X., Chen, Z., Wang, X., Dai, J., Wang, X., Huang, P., Zhao, Z., Shen, W.L., and Li, H. (2021). Periaqueductal gray neurons encode the sequential motor program in hunting behavior of mice. Nat. Commun. 12, 6523. 10.1038/s41467-021-26852-1.

35. Jesuthasan, S., Krishnan, S., Cheng, R.K., and Mathuru, A. (2021). Neural correlates of state transitions elicited by a chemosensory danger cue. Prog. Neuropsychopharmacol. Biol. Psychiatry 111. 10.1016/J.PNPBP.2020.110110.

36. Egan, R.J., Bergner, C.L., Hart, P.C., Cachat, J.M., Canavello, P.R., Elegante, M.F., Elkhayat, S.I., Bartels, B.K., Tien, A.K., Tien, D.H., et al. (2009). Understanding behavioral and physiological phenotypes of stress and anxiety in zebrafish. Behav. Brain Res. 205, 38–44. 10.1016/j.bbr.2009.06.022.

37. Agetsuma, M., Aizawa, H., Aoki, T., Nakayama, R., Takahoko, M., Goto, M., Sassa, T., Amo, R., Shiraki, T., Kawakami, K., et al. (2010). The habenula is crucial for experience-dependent modification of fear responses in zebrafish. Nat. Neurosci. 2010 1311 13, 1354–1356. 10.1038/nn.2654.

38. Signoret-Genest, J., Schukraft, N., L. Reis, S., Segebarth, D., Deisseroth, K., and Tovote, P. (2023). Integrated cardio-behavioral responses to threat define defensive states. Nat. Neurosci. 26, 447–457. 10.1038/s41593-022-01252-w.

39. Khan, B., Jaesiri, O., Lazarte, I.P., Li, Y., Tian, G., Zhao, P., Zhao, Y., Ho, V.D., and Semmelhack, J.L. (2023). Zebrafish larvae use stimulus intensity and contrast to estimate distance to prey. Curr. Biol. 10.1016/j.cub.2023.06.046.

40. Khan, B., Lazarte, I.P., Jaesiri, O., Zhao, P., and Semmelhack, J.L. (2024). Protocol for using UV stimuli to evoke prey capture strikes in head-fixed zebrafish larvae. STAR Protoc. 5, 102780. 10.1016/j.xpro.2023.102780.

41. Bianco, I.H., Kampff, A.R., and Engert, F. (2011). Prey capture behavior evoked by simple visual stimuli in larval zebrafish. Front. Syst. Neurosci. 5, 101. 10.3389/fnsys.2011.00101.

42. Freeman, J., Vladimirov, N., Kawashima, T., Mu, Y., Sofroniew, N.J., Bennett, D.V., Rosen, J., Yang, C.-T., Looger, L.L., and Ahrens, M.B. (2014). Mapping brain activity at scale with cluster computing. Nat. Methods 11, 941–950. 10.1038/nmeth.3041.

43. Pachitariu, M., Stringer, C., Dipoppa, M., Schröder, S., Rossi, L.F., Dalgleish, H., Carandini, M., and Harris, K.D. (2017). Suite2p: beyond 10,000 neurons with standard two-photon microscopy. bioRxiv, 061507. 10.1101/061507.

44. Kunst, M., Laurell, E., Mokayes, N., Kramer, A., Kubo, F., Fernandes, A.M., Förster, D., Dal Maschio, M., and Baier, H. (2019). A Cellular-Resolution Atlas of the Larval Zebrafish Brain. Neuron 103, 21–38.e5. 10.1016/j.neuron.2019.04.034.

45. Chen, X., Mu, Y., Hu, Y., Kuan, A.T., Nikitchenko, M., Randlett, O., Chen, A.B., Gavornik, J.P., Sompolinsky, H., Engert, F., et al. (2018). Brain-wide Organization of Neuronal Activity and Convergent Sensorimotor Transformations in Larval Zebrafish. Neuron 100, 876–890.e5. 10.1016/j.neuron.2018.09.042.

46. Marques, J.C., Li, M., Schaak, D., Robson, D.N., and Li, J.M. (2020). Internal state dynamics shape brainwide activity and foraging behaviour. Nature 577, 239–243. 10.1038/s41586-019-1858-z.

47. Bianco, I.H., and Engert, F. (2015). Visuomotor Transformations Underlying Hunting Behavior in Zebrafish. Curr. Biol. 25, 831–846. 10.1016/j.cub.2015.01.042.

48. Herrero, L., Rodríguez, F., Salas, C., and Torres, B. (1998). Tail and eye movements evoked by electrical microstimulation of the optic tectum in goldfish. Exp. Brain Res. 120, 291–305.

49. Helmbrecht, T.O., dal Maschio, M., Donovan, J.C., Koutsouli, S., and Baier, H. (2018). Topography of a Visuomotor Transformation. Neuron. 10.1016/j.neuron.2018.10.021.

50. Scott, E.K., Mason, L., Arrenberg, A.B., Ziv, L., Gosse, N.J., Xiao, T., Chi, N.C., Asakawa, K., Kawakami, K., and Baier, H. (2007). Targeting neural circuitry in zebrafish using GAL4 enhancer trapping. Nat. Methods 4, 323–326. 10.1038/nmeth1033.

51. Pearl, J. (2009). Causality 2nd ed. (Cambridge University Press) 10.1017/CBO9780511803161.

52. Mueller, T., Dong, Z., Berberoglu, M.A., and Guo, S. (2011). The dorsal pallium in zebrafish, Danio rerio (Cyprinidae, Teleostei). Brain Res. 1381, 95–105. 10.1016/j.brainres.2010.12.089.

53. Procacci, N.M., Allen, K.M., Robb, G.E., Ijekah, R., Lynam, H., and Hoy, J.L. (2020). Context-dependent modulation of natural approach behaviour in mice. Proc. R. Soc. B Biol. Sci. 287, 20201189. 10.1098/rspb.2020.1189.

54. Knudsen, E.I. (1982). Auditory and visual maps of space in the optic tectum of the owl. J. Neurosci. Off. J. Soc. Neurosci. 2, 1177–1194. 10.1523/JNEUROSCI.02-09-01177.1982.

55. Heap, L.A., Vanwalleghem, G.C., Thompson, A.W., Favre-Bulle, I., Rubinsztein-Dunlop, H., and Scott, E.K. (2018). Hypothalamic projections to the optic tectum in larval zebrafish. Front. Neuroanat. 11, 135.

56. Henriques, P.M., Rahman, N., Jackson, S.E., and Bianco, I.H. (2019). Nucleus Isthmi Is Required to Sustain Target Pursuit during Visually Guided Prey-Catching. Curr. Biol. CB 29, 1771–1786.e5. 10.1016/j.cub.2019.04.064.

57. Mysore, S.P., and Knudsen, E.I. (2011). The role of a midbrain network in competitive stimulus selection. Curr. Opin. Neurobiol. 21, 653–660. 10.1016/j.conb.2011.05.024.

58. Zaupa, M., Nagaraj, N., Sylenko, A., Baier, H., Sawamiphak, S., and Filosa, A. (2024). The Calmodulin-interacting peptide Pcp4a regulates feeding state-dependent behavioral choice in zebrafish. Neuron.

59. Yáñez, J., Suárez, T., Quelle Regaldie, A., Folgueira, M., and Anadon, R. (2018). Neural connections of the pretectum in zebrafish ( Danio rerio ). J. Comp. Neurol. 526. 10.1002/cne.24388.

60. Romano, S.A., Pietri, T., Pérez-Schuster, V., Jouary, A., Haudrechy, M., and Sumbre, G. (2015). Spontaneous neuronal network dynamics reveal circuit’s functional adaptations for behavior. Neuron 85, 1070–1085. 10.1016/j.neuron.2015.01.027.

61. Marquez-Legorreta, E., Constantin, L., Piber, M., Favre-Bulle, I.A., Taylor, M.A., Blevins, A.S., Giacomotto, J., Bassett, D.S., Vanwalleghem, G.C., and Scott, E.K. (2022). Brain-wide visual habituation networks in wild type and fmr1 zebrafish. Nat. Commun. 13, 895. 10.1038/s41467-022-28299-4.

62. Wei, P., Liu, N., Zhang, Z., Liu, X., Tang, Y., He, X., Wu, B., Zhou, Z., Liu, Y., Li, J., et al. (2015). Processing of visually evoked innate fear by a non-canonical thalamic pathway. Nat. Commun. 6, 6756. 10.1038/ncomms7756.

63. Lovett-Barron, M., Chen, R., Bradbury, S., Andalman, A.S., Wagle, M., Guo, S., and Deisseroth, K. (2020). Multiple convergent hypothalamus–brainstem circuits drive defensive behavior. Nat. Neurosci. 23, 959–967. 10.1038/s41593-020-0655-1.

64. Marquart, G.D., Tabor, K.M., Bergeron, S.A., Briggman, K.L., and Burgess, H.A. (2019). Prepontine non-giant neurons drive flexible escape behavior in zebrafish. PLOS Biol. 17, e3000480. 10.1371/journal.pbio.3000480.

65. Dana, H., Sun, Y., Mohar, B., Hulse, B.K., Kerlin, A.M., Hasseman, J.P., Tsegaye, G., Tsang, A., Wong, A., Patel, R., et al. (2019). High-performance calcium sensors for imaging activity in neuronal populations and microcompartments. Nat. Methods 16, 649–657. 10.1038/s41592-019-0435-6.

