## Supplemental Figures and Tables for "The visuomotor transformations underlying target-directed behavior"

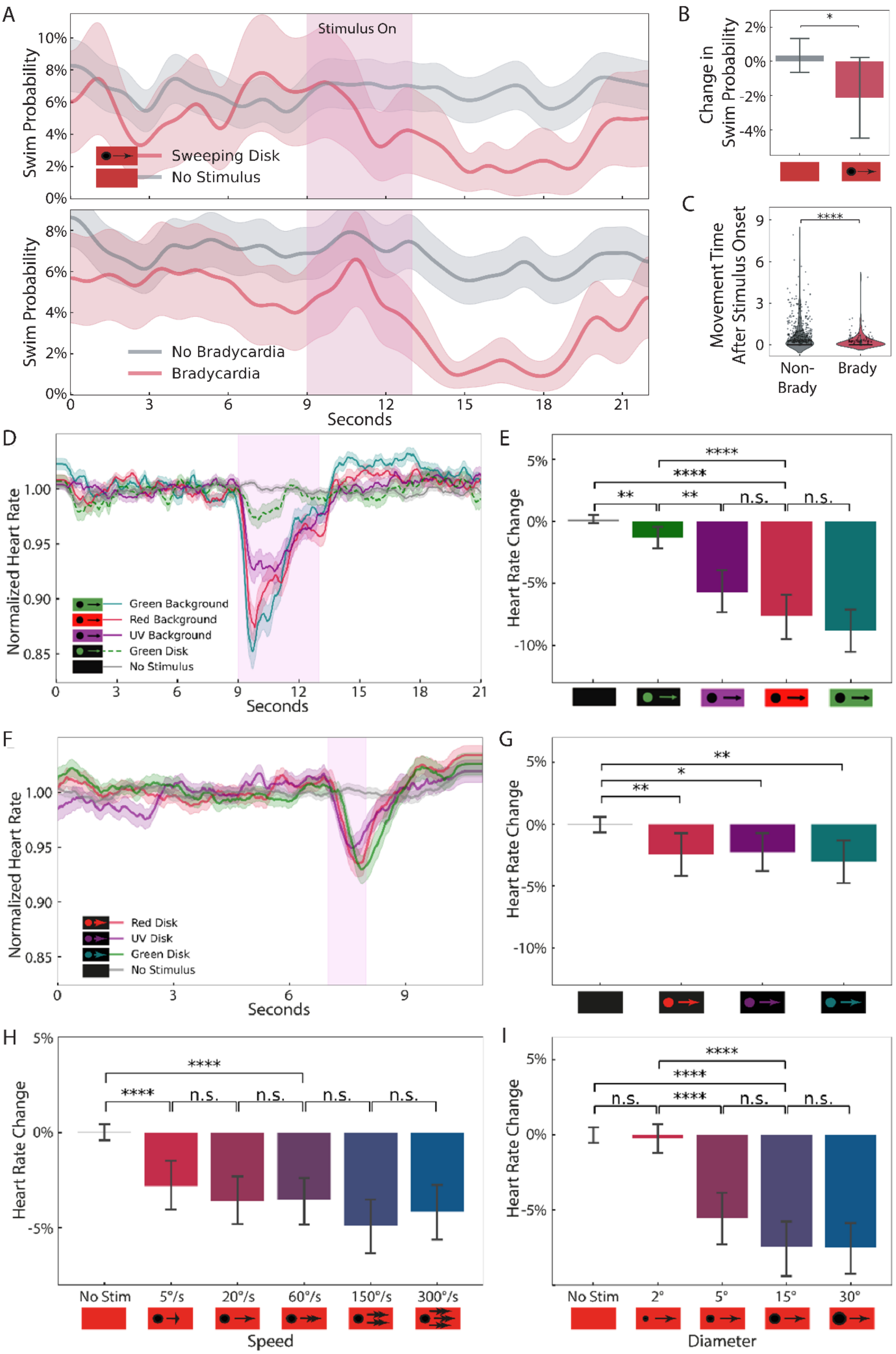
A
Stimulus On
Swim Probability
Sweeping Disk
No Stimulus
No Bradycardia
Bradycardia
Seconds
B
Change in Swim Probability
C
Movement Time After Stimulus Onset
Non-Brady
Brady
D
Normalized Heart Rate
Green Background
Red Background
UV Background
Green Disk
No Stimulus
E
Heart Rate Change
F
Red Disk
UV Disk
Green Disk
No Stimulus
G
H
No Stim
5°/s
20°/s
60°/s
150°/s
300°/s
Speed
I
2°
5°
15°
30°
Diameter

**Supplementary Figure 1: Characterization of the freezing stimulus parameters.** (A) Normalized heart rate over time in response to sweeping disks with different stimulus and background colors. Red background and no stimulus data are the same as in figure 1E. Pink bar indicates 4-second stimulus presentation. Shading shows standard error (n = 14). (B) Change in heart rate during the 4-second stimulus window. (C) Normalized heart rate over time in response to a sweeping disks of different colors on a dark background. Pink bar indicates 1-second stimulus presentation (n = 13). (D) Change in heart rate during the 3 seconds after stimulus onset for stimuli in C. (E) Heart rate change during the 4-second stimulus window caused by sweeping disks of different diameters. (F) Heart rate change during the same window for sweeping disks of varying speed (n = 11). *, $p<0.05$. **, $p<0.01$. ****, $p<0.0001$, Kruskal-Wallis test, followed by Dunn's test. Error bar represents standard deviation.

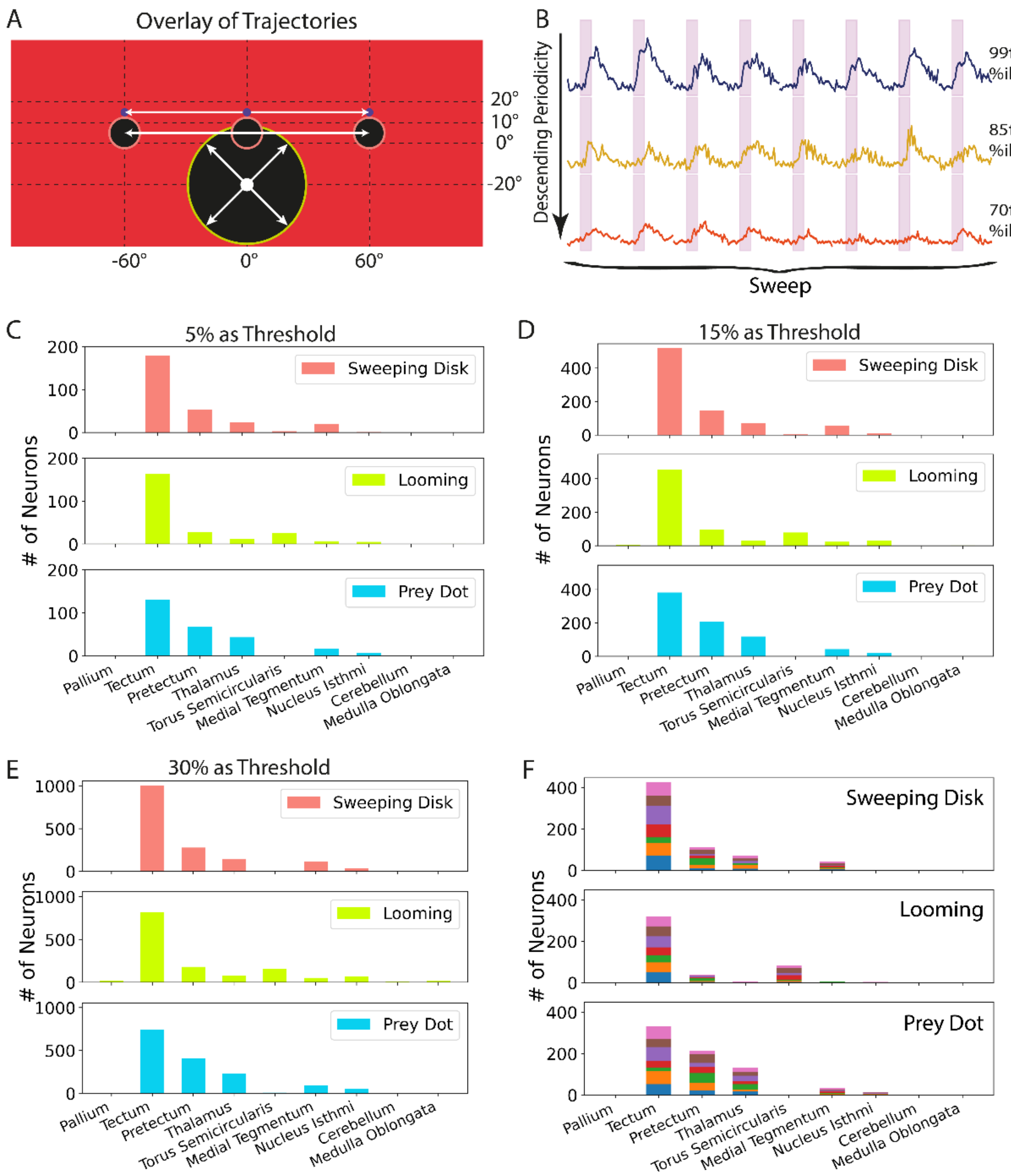


**Supplementary Figure 2: Visual stimuli and setting the threshold for sensory neurons.** (A) Schematic representing the movement of each stimulus and its relative position on the screen. (B) Example ΔF/F traces during 8 presentations of the sweep stimulus (pink shading) for neurons with sensory indices from the $99^{th}$, $85^{th}$, and $70^{th}$ percentile of the sweep sensory index. (C-E) The distrbution of sensory neurons in each brain area based on using the top 10% most correlated neurons and then a threshold of the top 5, 15, or 30% of the neurons by sensory index, as compared to the threshold of 15% used in figure 3. No spatial colocalization filtering was applied for this analysis. (F) The anatomical distribution of each type of sensory neuron across the seven fish, using the top 15% of SI as the threshold. Each color represents the sensory neurons from one fish.

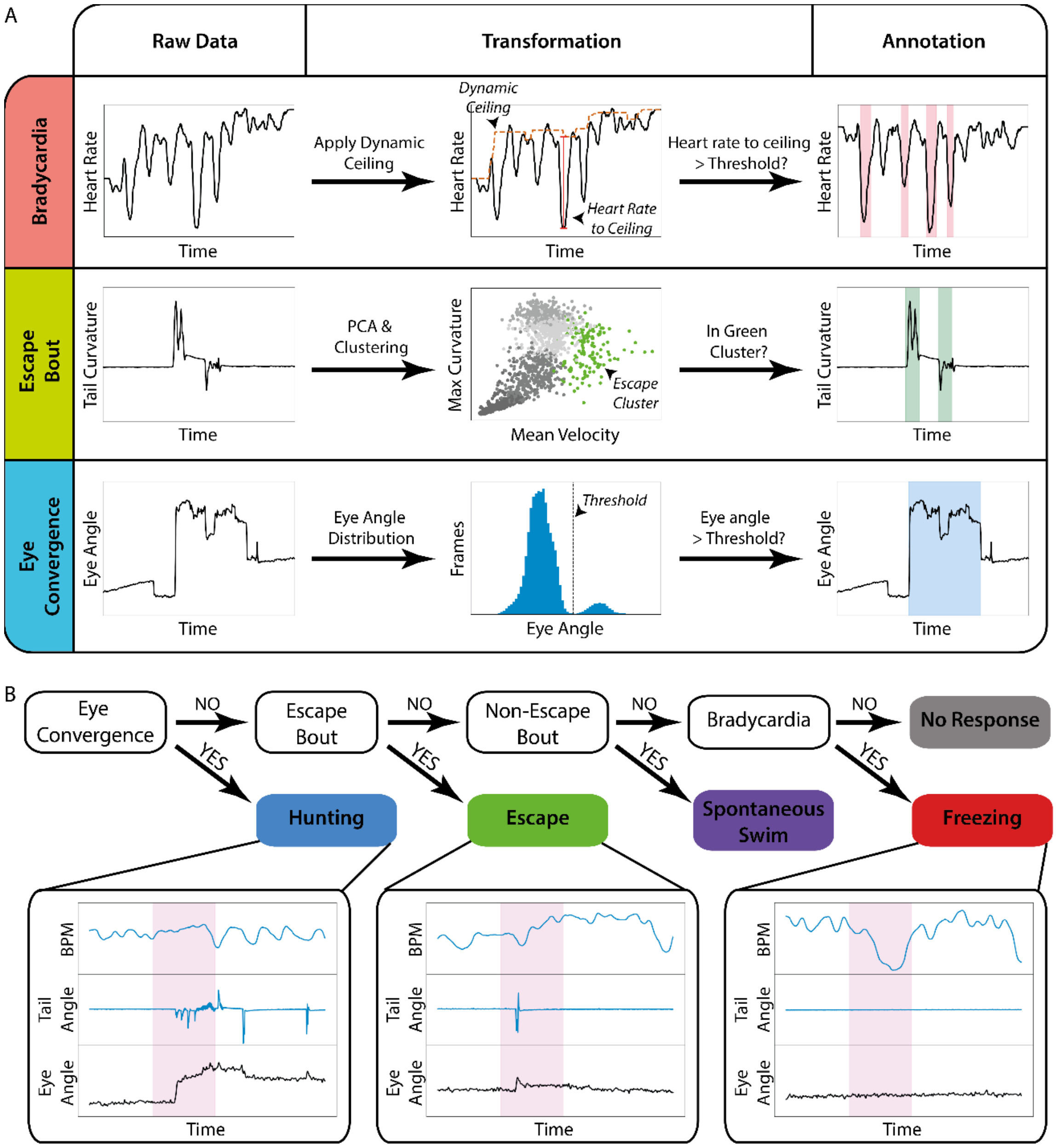


**Supplementary Figure 3: The categorization of behavioral responses.** (A) Annotation of bradycardia, escape, and eye convergence. Frames were annotated as bradycardia if the heart-rate-to-ceiling distance to ceiling is larger than 3 standard deviations of the heart-rate-to-ceiling distance during no stimulus trials. For escape, any bout belonging to the cluster with high tail velocity and curvature (green) was annotated as an escape bout. To find eye convergence bouts, the distribution of eye angles from each animal was plotted, and the first local minimum larger than the maximum in the distribution was set as the threshold for eye convergence. Any frame with eye angle higher than the threshold was annotated as eye convergence. (B) The sequential pipleline for classification. Examples of each behavior are shown below, with corresponding heart, tail and eye angle data.

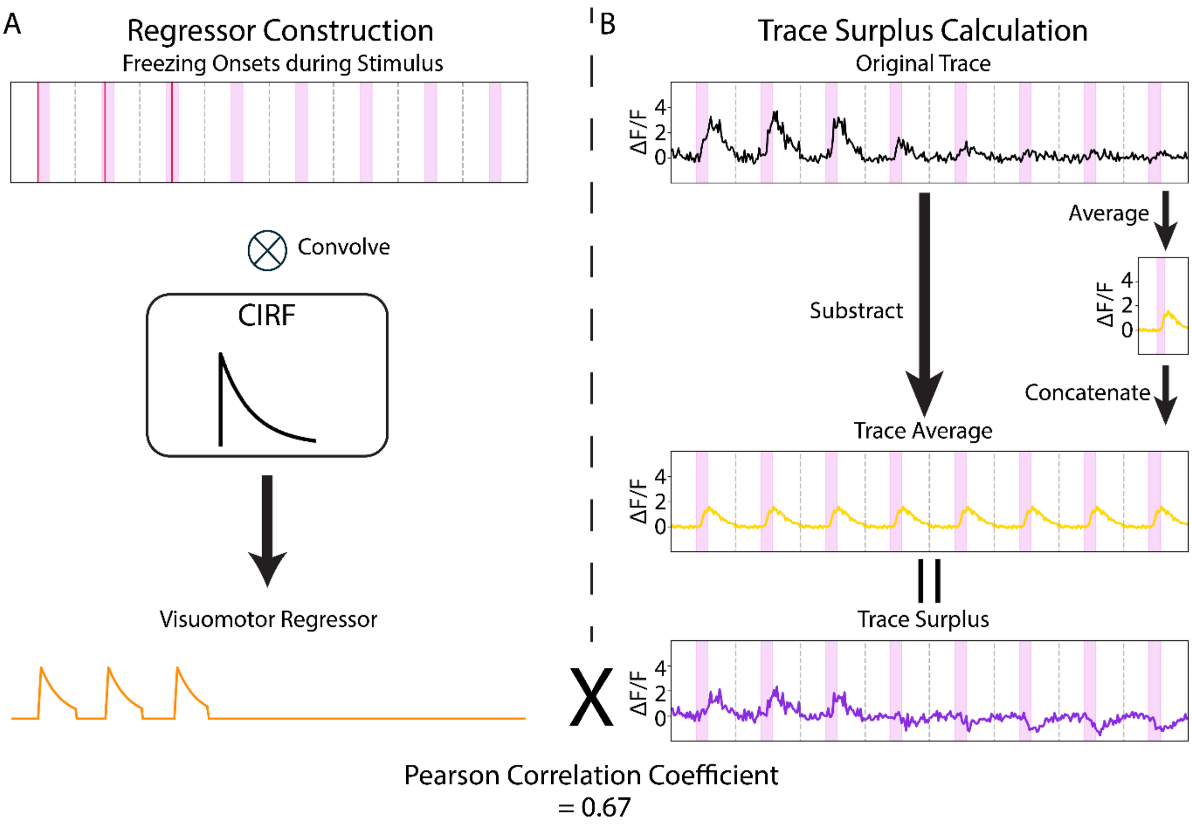


**Supplementary Figure 4: Construction of the visuomotor regressors and calculation of correlation score.** (A) Example freezing behavior for one fish in the imaging dataset. The freezing regressor was constructed based on the onset of each freezing event (bradycardia + no movement) during the stimulus time period (red lines). The behavior onset time was shifted by -.5 seconds so that the response starts before the behavior, and convolved with a Calcium Impulse Response Function (CIRF) to generate the visuomotor regressor. Regressors for hunting and escape were constructed similarly, using the initiation of eye convergence or the escape bout as behavior onset. (B) Example freezing SM neuron trace from our dataset. The trace average was subtracted from the trace to generate a trace surplus. For this neuron, the Pearson Correlation between the visuomotor regressor and the trace surplus was 0.67.

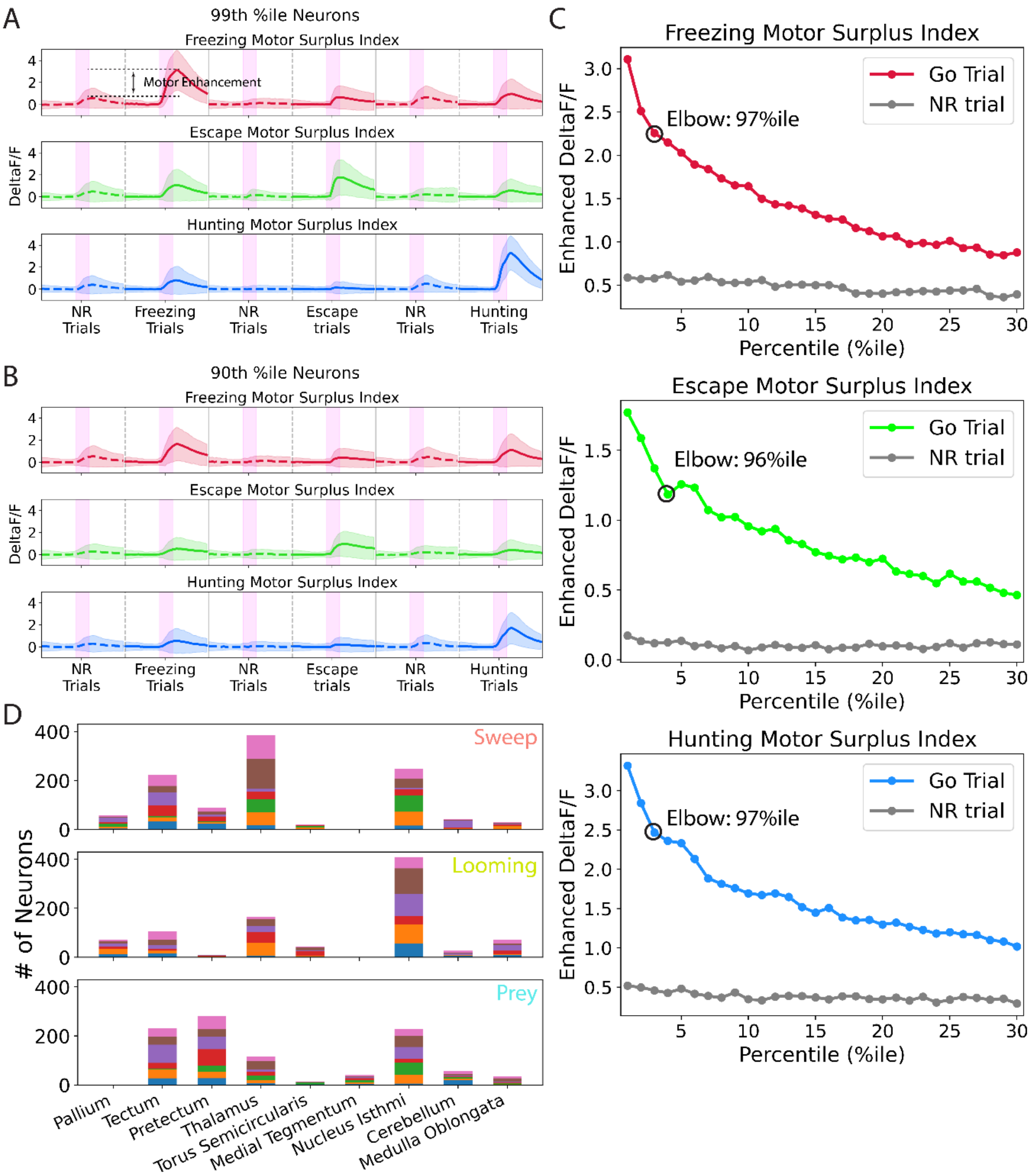


**Supplementary Figure 5: Setting the threshold for sensorimotor neurons.** (A-B) Average calcium response in behavior and no response trials for sensory-motor neurons from the 99$^{th}$ (A) and 90$^{th}$ (B) percentile of MSI. Motor enhancement is the difference between the peak ΔF/F in response and no response trials. Pink bars represent 4-second presentation of sweep, prey or looming stimuli. n = 7 larvae. (C) Motor enhancement (colored line) for each percentile of neurons on the freezing, escape, and hunting MSI. An elbow was located near the 97$^{th}$ percentile for all three distributions, and this was set as the threshold for SM neurons. Grey line represents peak ΔF/F of that percentile of neurons in NR trials. (D) Anatomical distribution of SM neurons across the seven fish; each color represents SM neurons of that type in one fish. One fish was excluded from the looming SM neuron calculation due to its having no trials without behavior.

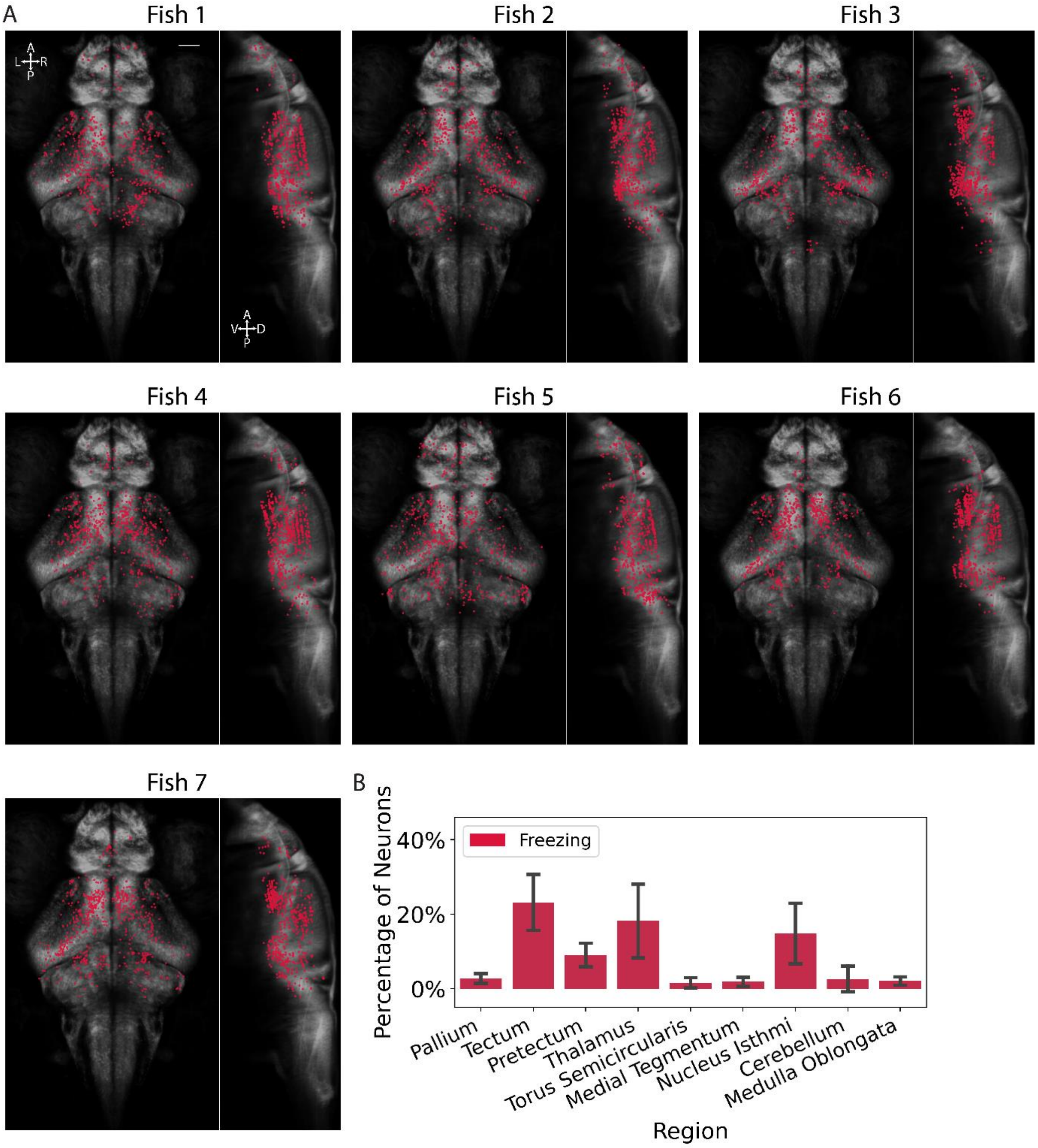


**Supplementary Figure 6: Neurons with high freezing MSI are found in the same anatomical regions across seven larvae.** (A) Anatomical locations of the top 600 neurons by freezing MSI in each fish, without spatial colocalization filtering. (B) Percentage of the top 600 freezing MSI neurons in each area across seven fish. Error bar represents Standard Deviation.

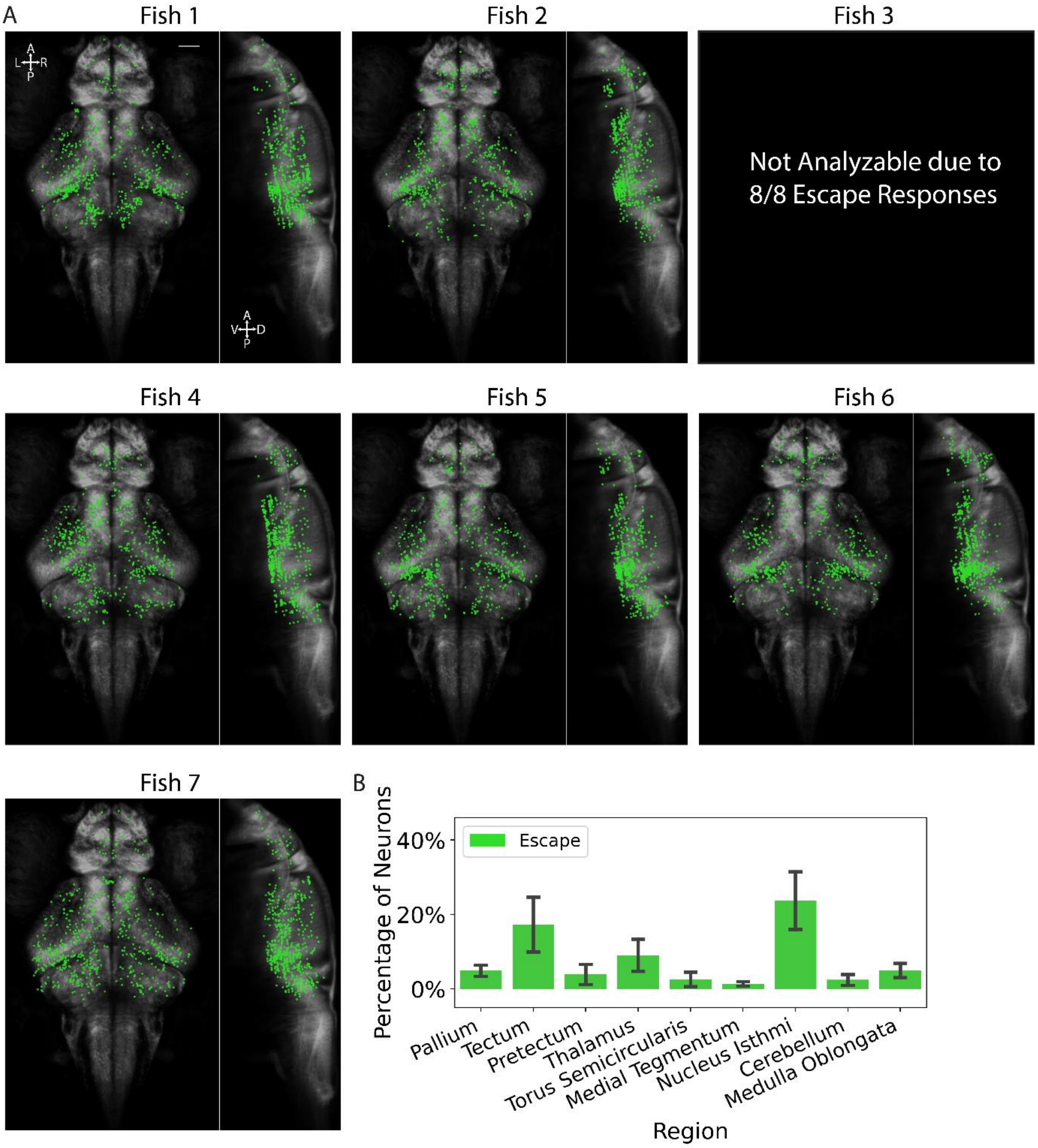


**Supplementary Figure 7: Neurons with high escape MSI are found in the same anatomical regions across six larvae.** (A) Anatomical locations of the top 600 neurons by escape MSI in each fish, without spatial colocalization filtering. (B) Percentage of top 600 escape MSI neurons in each area across seven fish. Error bar represents Standard Deviation.

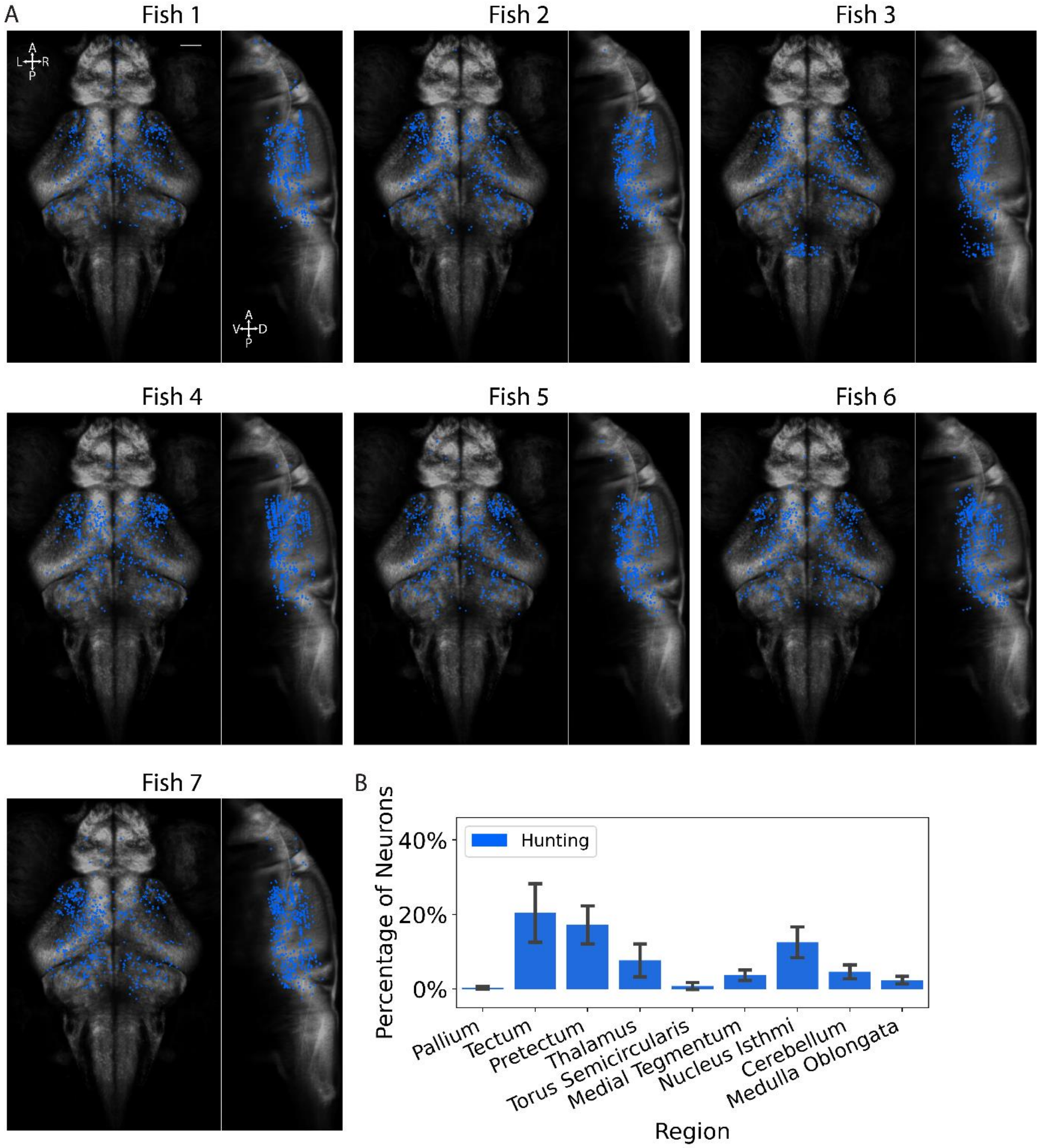


**Supplementary Figure 8: Neurons with high hunting MSI are found in the same anatomical regions across seven larvae.** (A) Anatomical locations of the top 600 neurons by hunting MSI in each fish, without spatial colocalization filtering. (B) Percentage of the top 600 hunting MSI neurons in each area across seven fish. Error bar represents Standard Deviation.

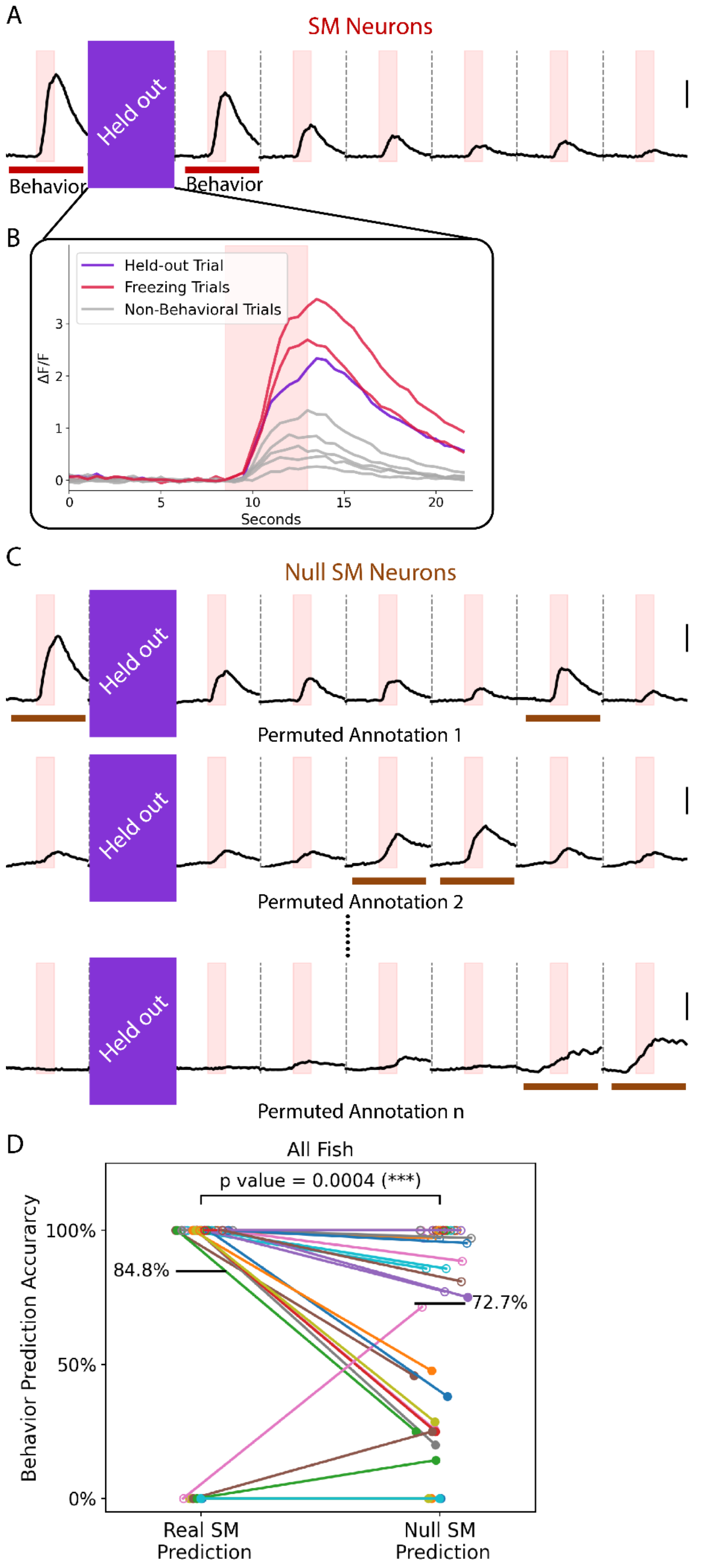


**Supplementary Figure 9: Freezing SM neurons can predict freezing behavior.** (A) Average response of the freezing SM neurons in one fish, selected based on activity in 7 trials, with the second trial held out. Scale bar represents ΔF/F = 1. (B) Activity during the held-out trial (purple) compared to behavior (red) and no response trials (grey). The maximum value of the held-out trace was compared to the same value for behavior and non-behavior trials to make the prediction. (C) Average activity of the null SM neurons selected based on different permutations of the freezing annotation. Brown bars represent the freezing trials for each permutation. (D) Accuracy with which SM neurons predict the behavioral annotation (freezing or no response) of the held out trial. For null SM neurons, the ability to predict the permuted annotation (brown bars) was tested. Each dot represents one held out trial in one fish. Solid dots represent freezing trials, and open dots represent no response trials. For the null SM neurons, the dots represent their accuracy over all possible permutations of freezing annotation with a given trial held out. ***, $p<0.001$, one sided Wilcoxon signed-rank test.

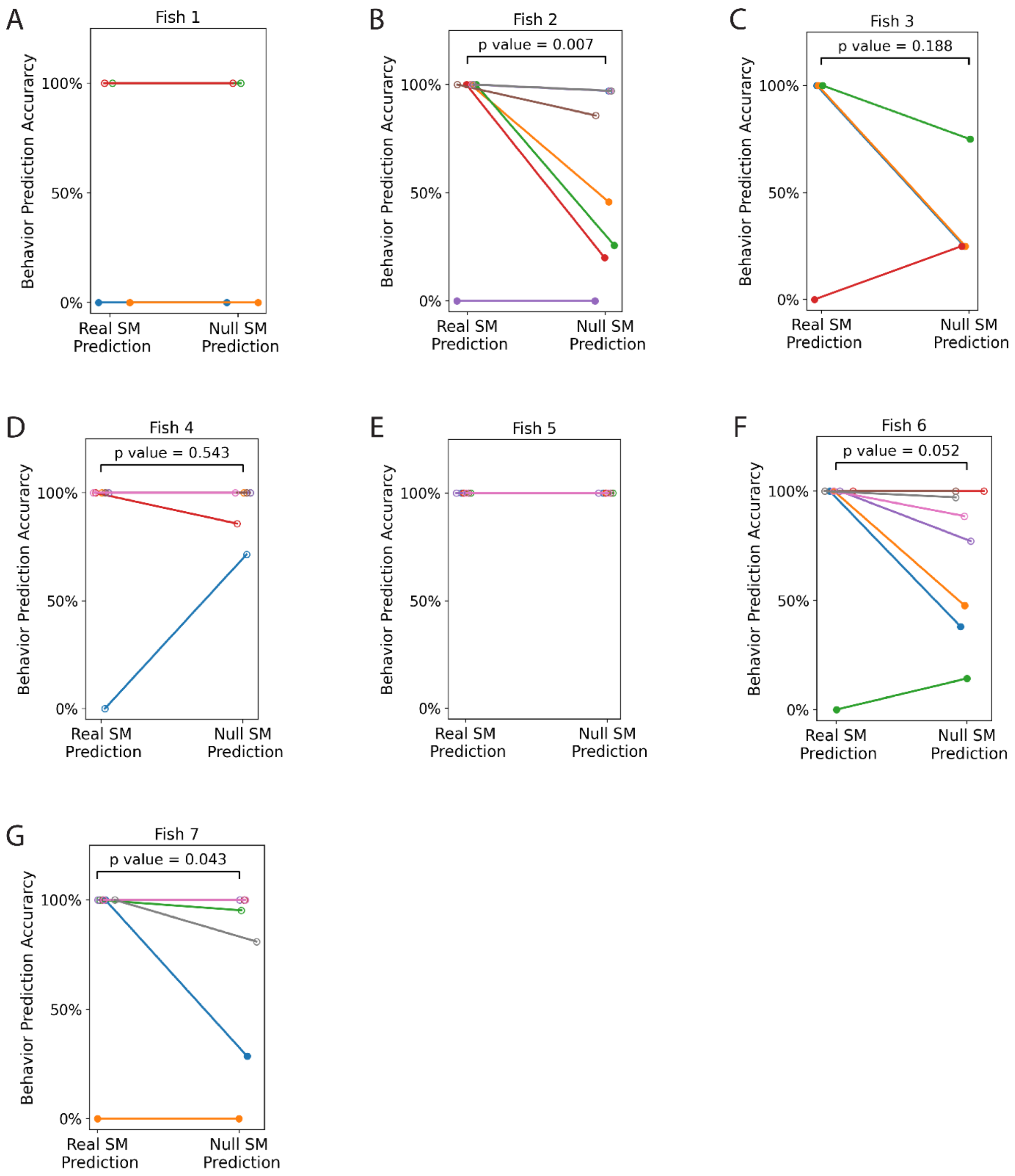


**Supplementary Figure 10: Behavioral prediction accuracy of real and null SM neurons in each fish.** (A-G) Accuracy with which SM neurons predict the behavioral annotation (freezing or no response) of the held out trial, as in Supplementary figure x, for each fish. Each dot represents one held out trial in one fish. Solid dots represent freezing trials, and open dots represent no response trials. P value is calculated with the one sided Wilcoxon signed-rank test.

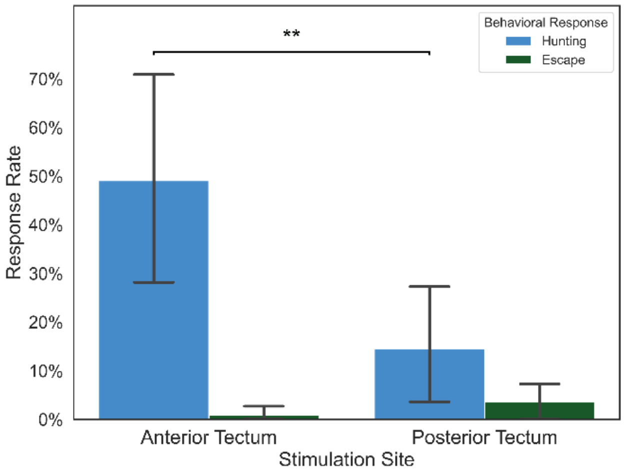


**Supplementary Figure 11: Hunting and escape responses triggered by optogenetic stimulation of Anterior or Posterior tectum.** Response rate is the percentage of trials annotated as having at least one episode of the behavior. Error bars represent Standard Deviation. n = 11 larvae. **, p<0.01. Wilcoxon Signed Rank test, one sided.

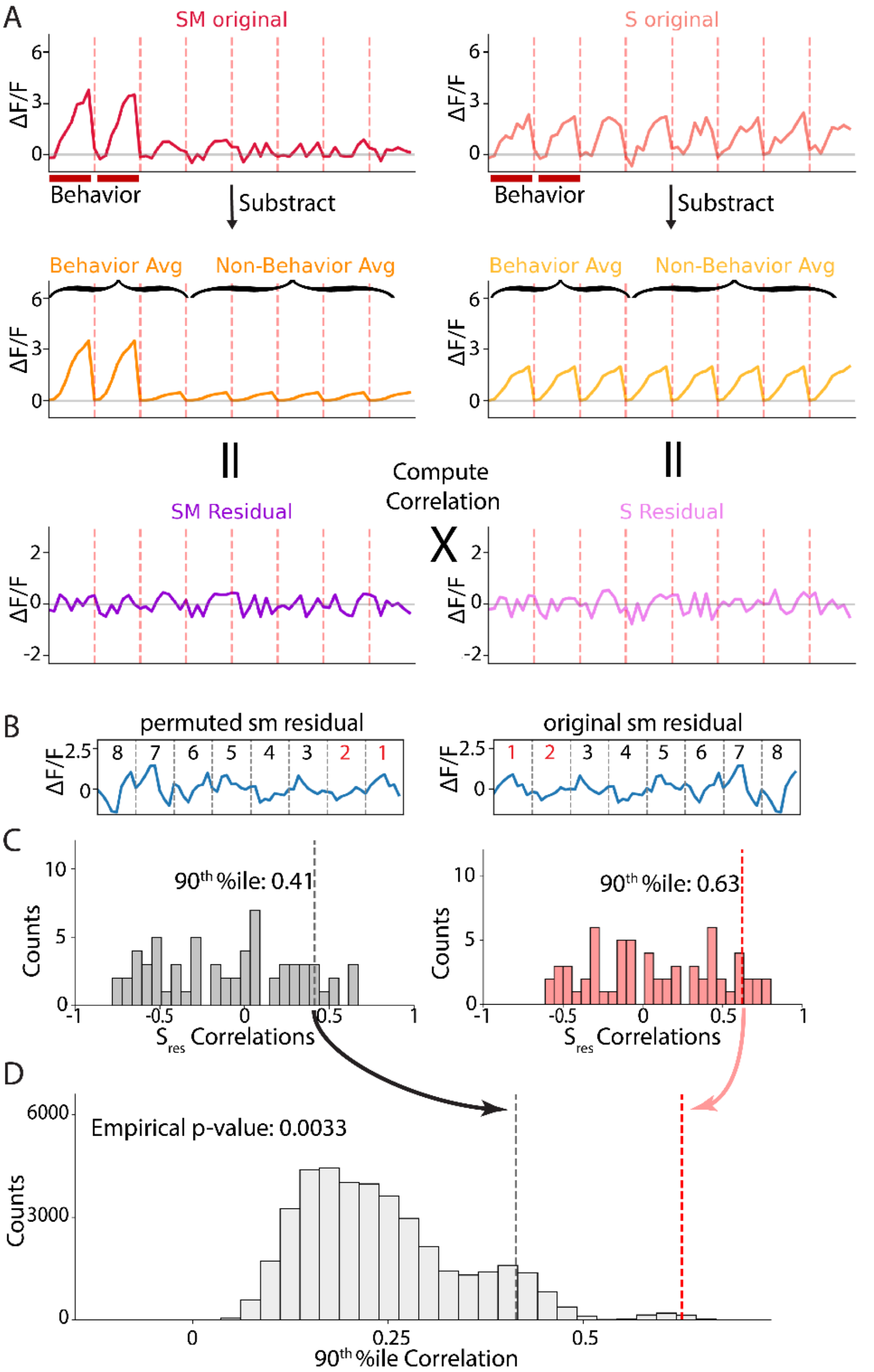


**Supplementary Figure 12: Calculation of p values for SM and S partial correlations.** (A) Calculation of SM and S residual traces and their correlation ρ(S,SM│B), i.e., the partial correlation, for a schematic freezing SM and sweep sensory neuron. (B) Example of trial-permuted and original SM residual traces for one freezing SM neuron. (C) Grey histogram: distribution of correlations between the permuted SM residual and the sweep sensory neurons in the same tectum. Pink histogram: distribution of correlations between the original SM residual and the sweep sensory neurons in the same tectum. Number of datapoints = Number of tectal sweep sensory neurons for that fish. (D) Grey dotted line: $90^{th}$%ile of one example permuted $SM_{res}$ to $S_{res}$ correlations. Light grey histogram: the distribution of the aforementioned $90^{th}$%ile's across the 8!=40,320 permutations of the SM residual. Red dotted line: $90^{th}$%ile of the original $S_{res}$ correlation with sensory residuals. Empirical p-value was calculated as the number of permutations whose $90^{th}$%ile correlation value was equal or greater than the original $SM_{res}$ $90^{th}$%ile correlation value (red dotted line) divided by the total number of permutations.

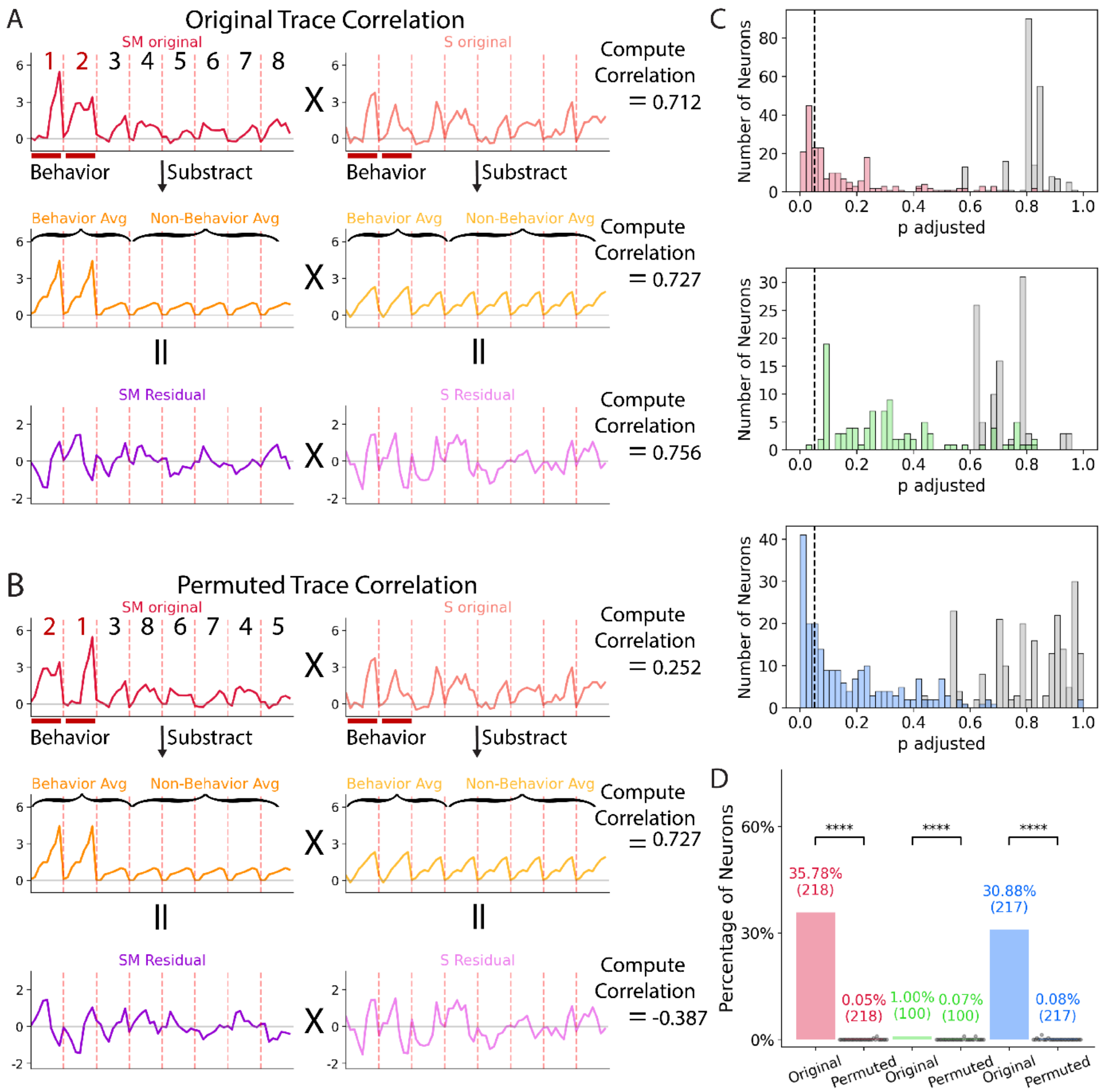


**Supplementary Figure 13: Subtraction of average traces does not create the correlation residuals.** (A) Calculation of SM to S residual correlation for a pair of example neurons from our dataset. (B) Calculation of the correlation between the S residual and the SM residual after the SM neuron's trace was permuted within behavior and non-behavior trials, leaving the behavior and non-behavior averages (middle row) unchanged. (C) Adjusted p-values for all freezing (red), escape (green), and hunting (blue) SM neurons, calculated by applying the Benjamini-Hochberg procedure to the empirical p-value from Supplementary figure 12, to adjust for testing all SM neurons combined, from all fish. Grey bars represent the adjusted p-values for SM neurons for one within category permutation as in B. (D) Percentage of each class of SM neuron that had a significant correlation with the corresponding S population in the original and permuted cases. In parenthesis is the total number of SM neurons of that type across all fish. Each black dot represents one of the 30 permutations sampled. ****, p<0.0001, One sided Wilcoxon signed-rank test.

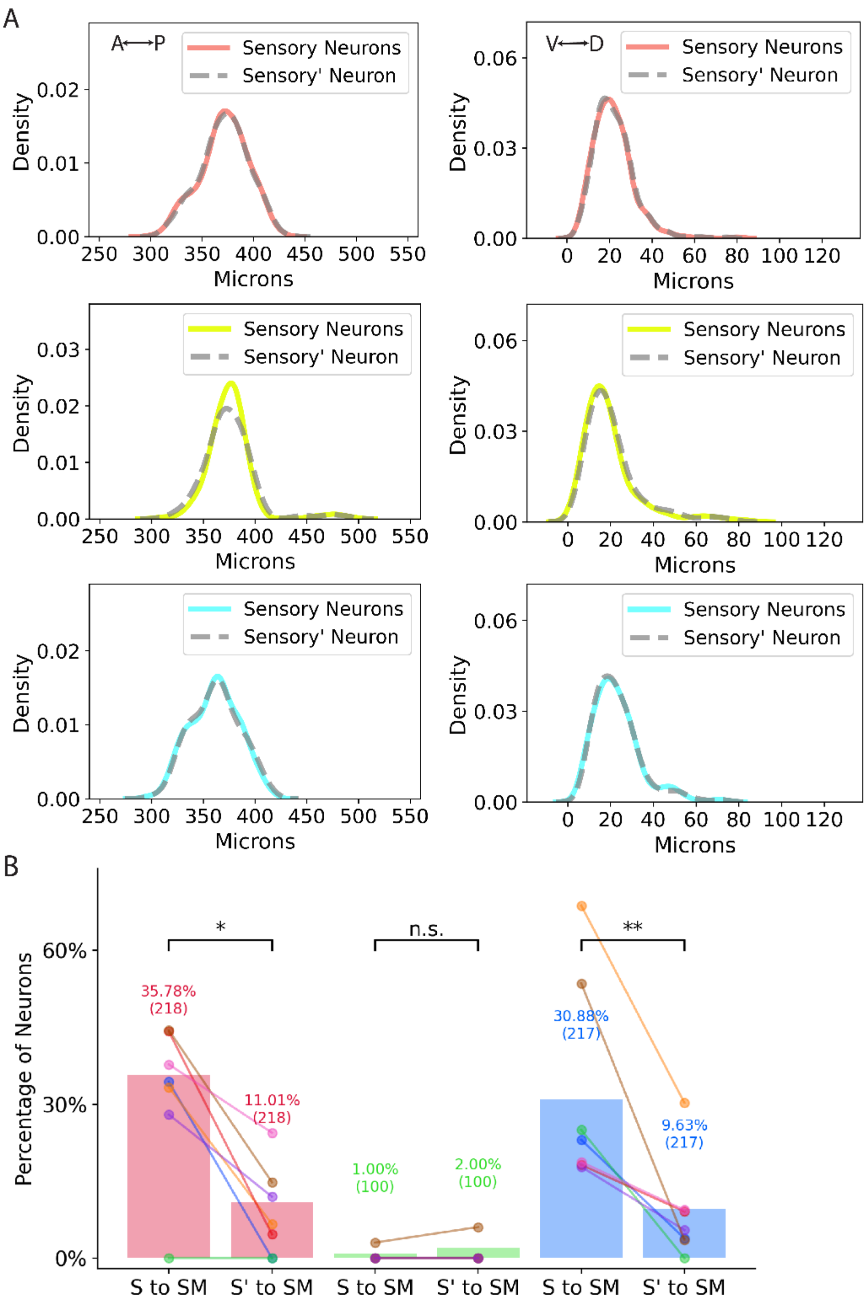


**Supplementary Figure 14: Anatomically matched surrogate S neurons with average SI indices are less correlated with SM neurons than the real sensory neurons.** (A) Location along the A/P and D/V axes of the tectum for sweep (red), looming (yellow-green), and prey (blue) sensory neurons and their S' counterparts (grey dashed lines). For each S neuron, an S' neuron was selected as the closest neuron in the same imaging plane with an SI +/-0.05 the average SI for tectal neurons in that fish (~.5) (B) Percentage of SM neurons that had significant correlations with S or S' neurons in the same tectum. In parenthesis is the total number of SM neurons of that type across all fish. Colored dots indicate the percentage of significant SM neurons in one fish. *, p<0.05; **, p<0.01, One sided Wilcoxon signed-rank test.

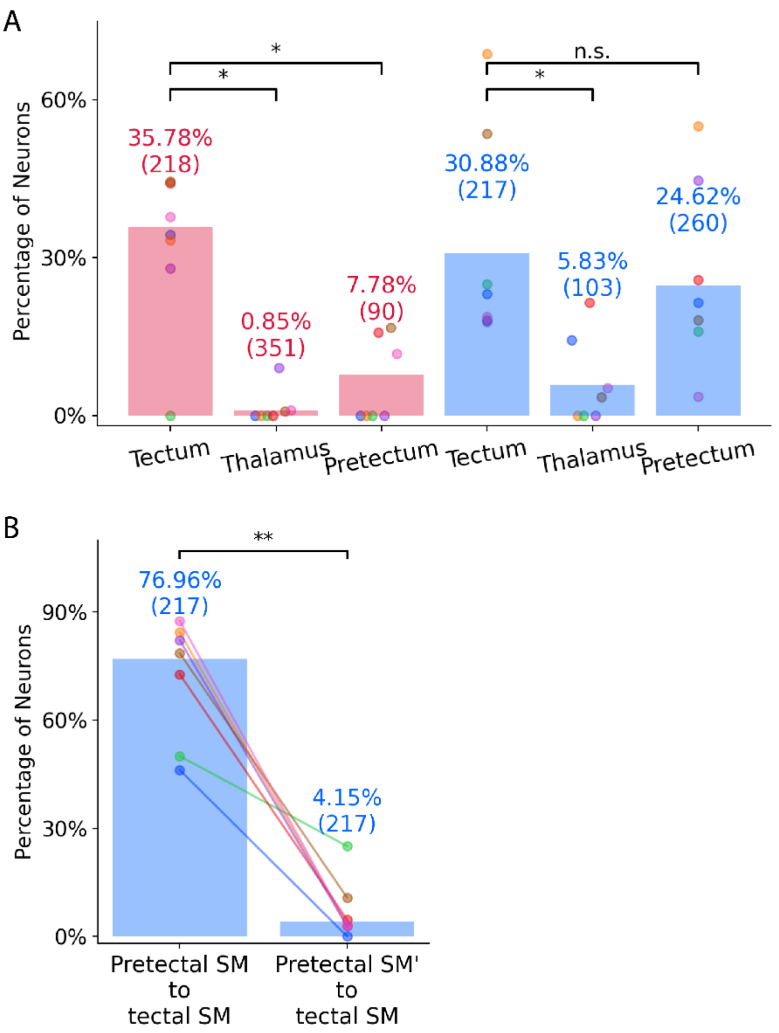


**Supplementary figure 15: Partial correlations in the tectum, pretectum, and thalamus.** (A) Bars represent the percentage of freezing SM (red) and hunting SM (blue) neurons in each area with a significant partial correlation with the S neurons in the same area. Each colored dot represents the number of significantly correlated SM neurons in that area in one fish. In parenthesis is the total number of SM neurons of that type across all fish. (B) Percentage of pretectal hunting SM neurons and SM' neurons with significant correlations to the tectal hunting SM population. For each pretectal SM neuron, an SM' neuron in the same imaging plane with an MSI +/-0.05 the average pretectal MSI for that fish was selected. *, $p<0.05$; **, $p<0.01$, One sided Wilcoxon signed-rank test.

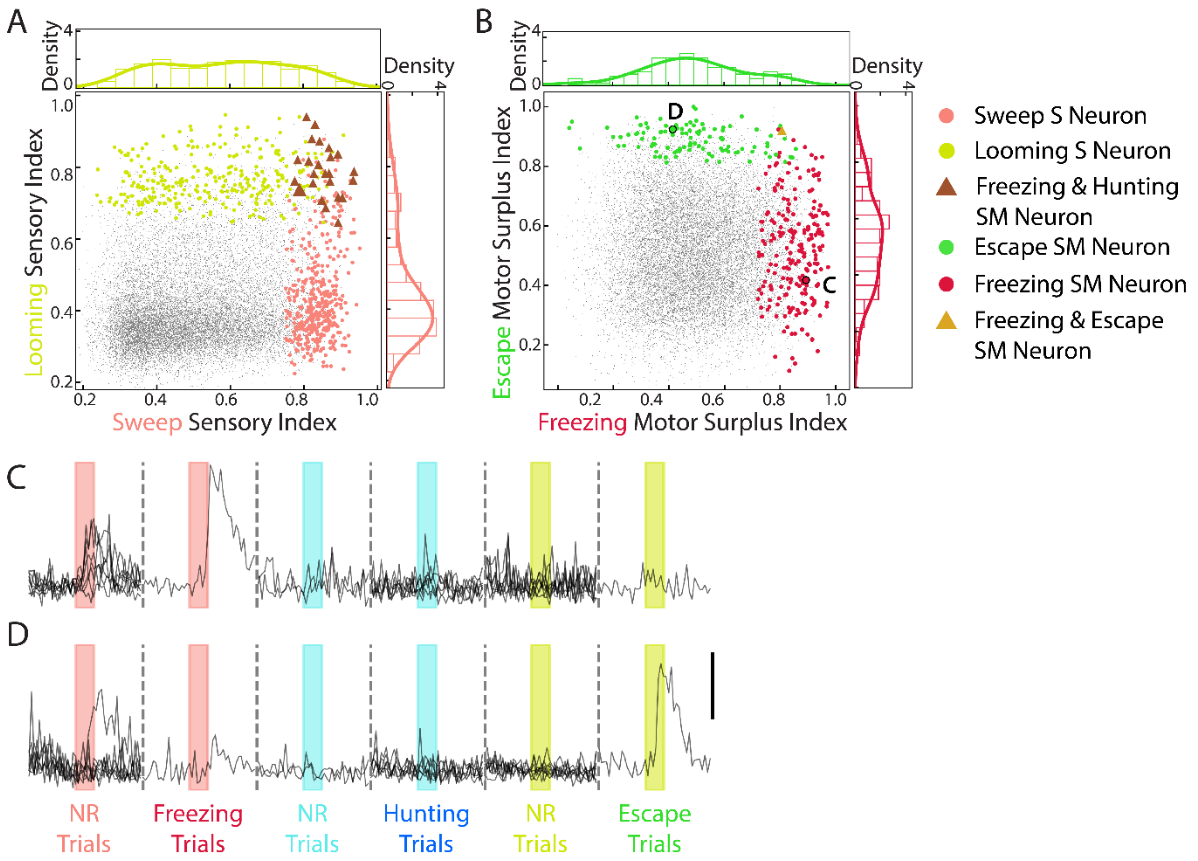


**Supplementary Figure 16: Functional segregation of defensive visual pathways in the tectum.** (A) The distribution of looming sensory neurons (light green) and sweep sensory neurons (light red) in the tectum on the looming and sweep SI. Crimson triangles: neurons belonging to both populations. (B) The distribution of escape (dark green) and freezing (dark red) SM neurons.Gold triangle = neuron belonging to both pobulations. (C-D) Responses of example freezing and escape SM neurons. Red, blue and green bars represent 4-second presentation of sweep, prey, and looming stimuli. Scale bar indicates ΔF/F = 3. The example neurons are noted in panel B.

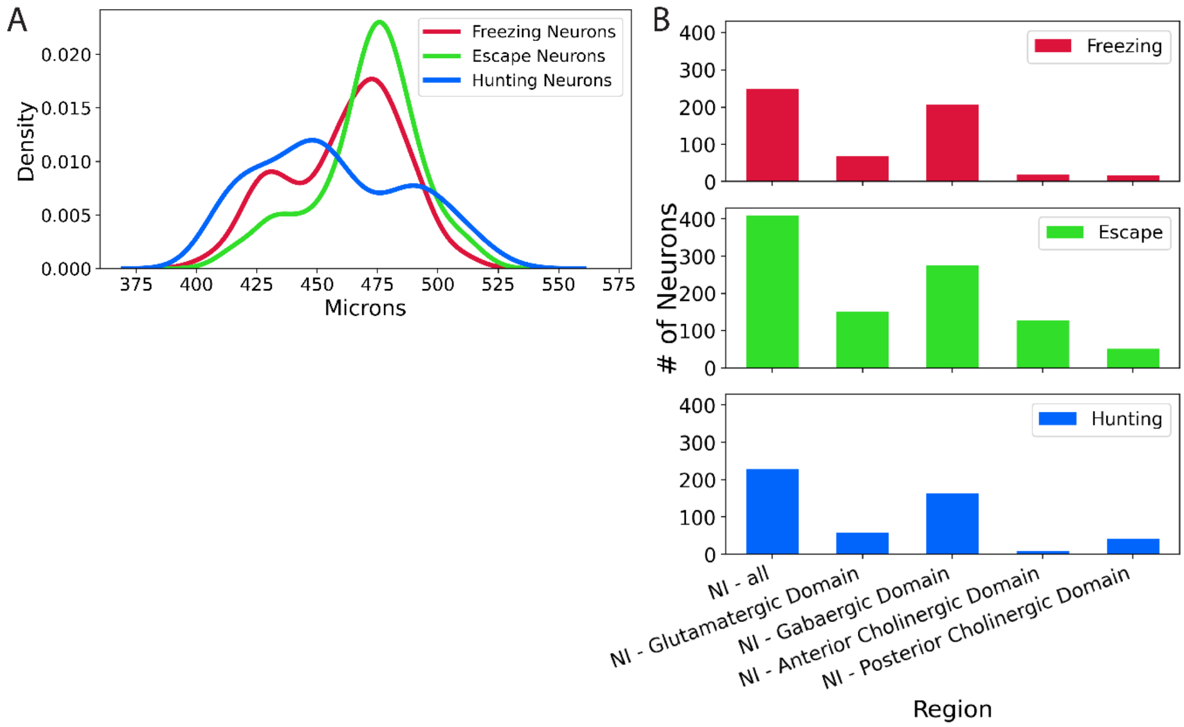


**Supplementary Figure 17: Anatomical distribution of SM neurons in the nucleus isthmi, and functional properties of all SM neurons.** (A) Distribution of hunting, freezing and escape SM neurons in the nucleus isthmi along the anterior to posterior axis. (B) Numbers of SM neurons total and in each subregion of the nucleus isthmi. (C) Average activity of SM neurons in the whole imaging volume during behavior or no response trials. Pink bars respresent 4-second stimulus presentation. n = 7 larvae.

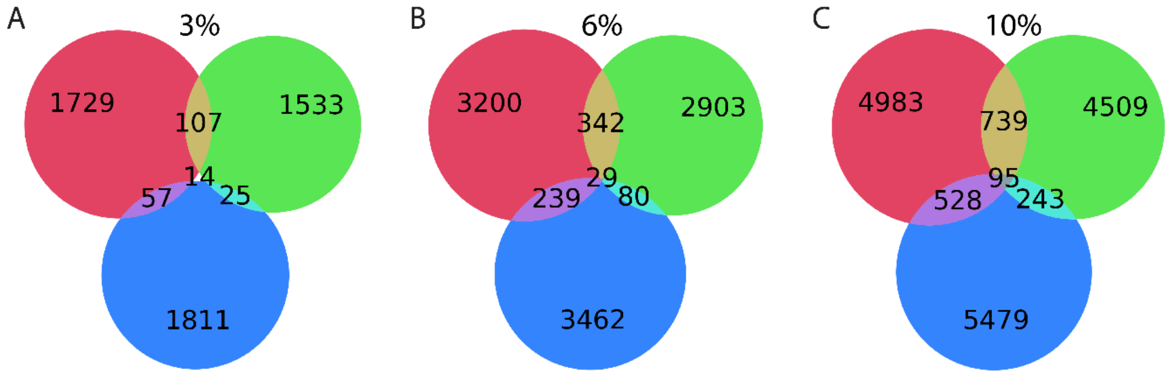


**Supplementary Figure 18: Functional properties of all SM neurons.** (A) Venn diagram showing the overlap among freezing (red), escape (green), and hunting (blue) SM populations using 3% as the threshiold. (B) Overlap when 6% is taken as the SM neuron threshold. (C) Overlap when 10% is taken as the SM neuron threshold.

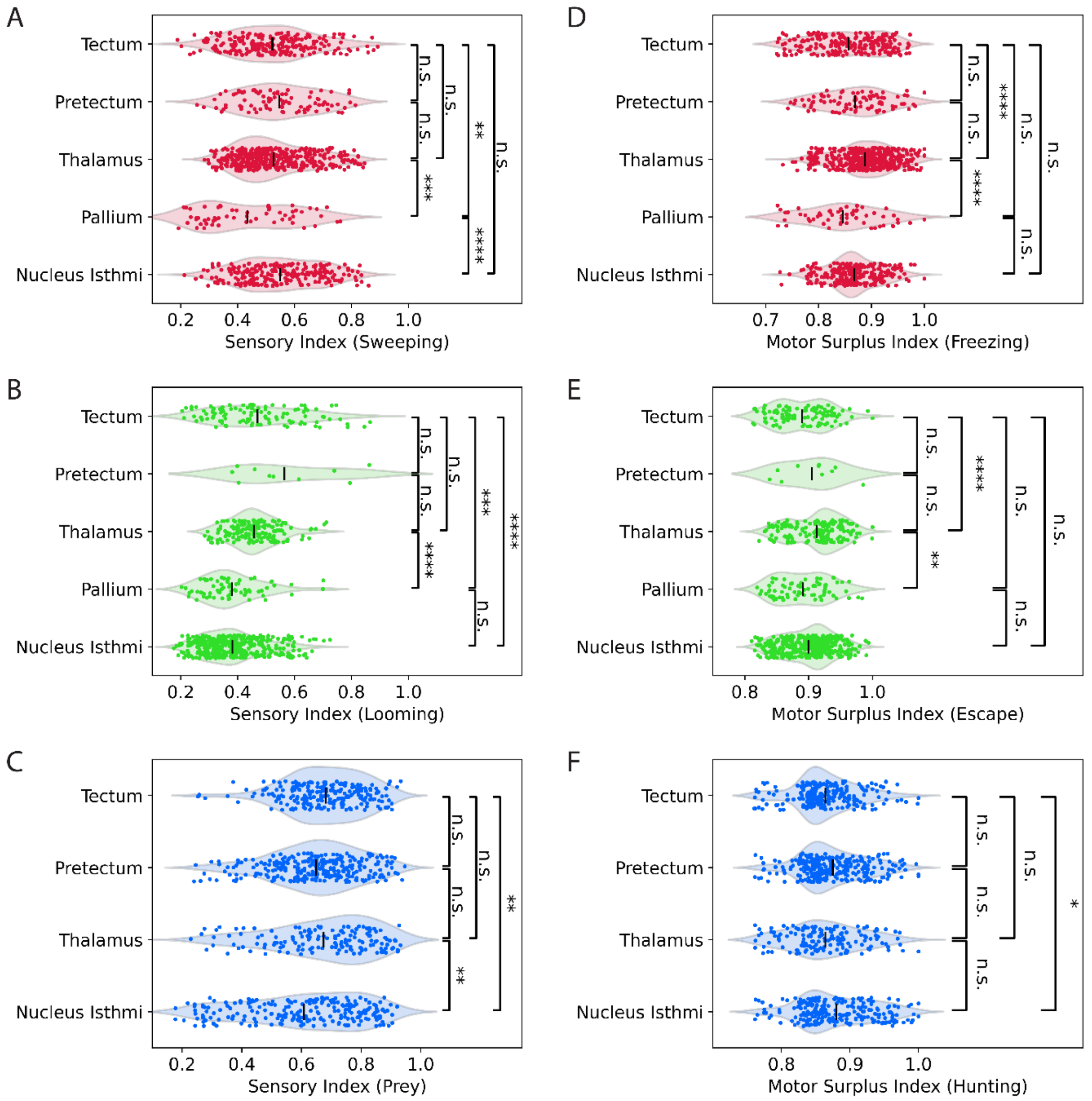


**Supplementary Figure 19: Sensory and visuomotor tuning of SM neurons in five brain regions.** (A) The sweep SI of freezing SM neurons in the five regions. (B) The looming SI of escape SM neurons. (C) The prey SI of hunting SM neurons. (D) The freezing MSI of freezing SM neurons in the five regions. (E) The escape MSI of escape SM neurons. (F) The hunting MSI of hunting SM neurons in the responsive brain areas. *, p<0.05. **, p<0.01. ***, p<0.001 ****, p<0.0001, Kruskal-Wallis test, followed by Dunn's test. Black lines indicate the mean value.

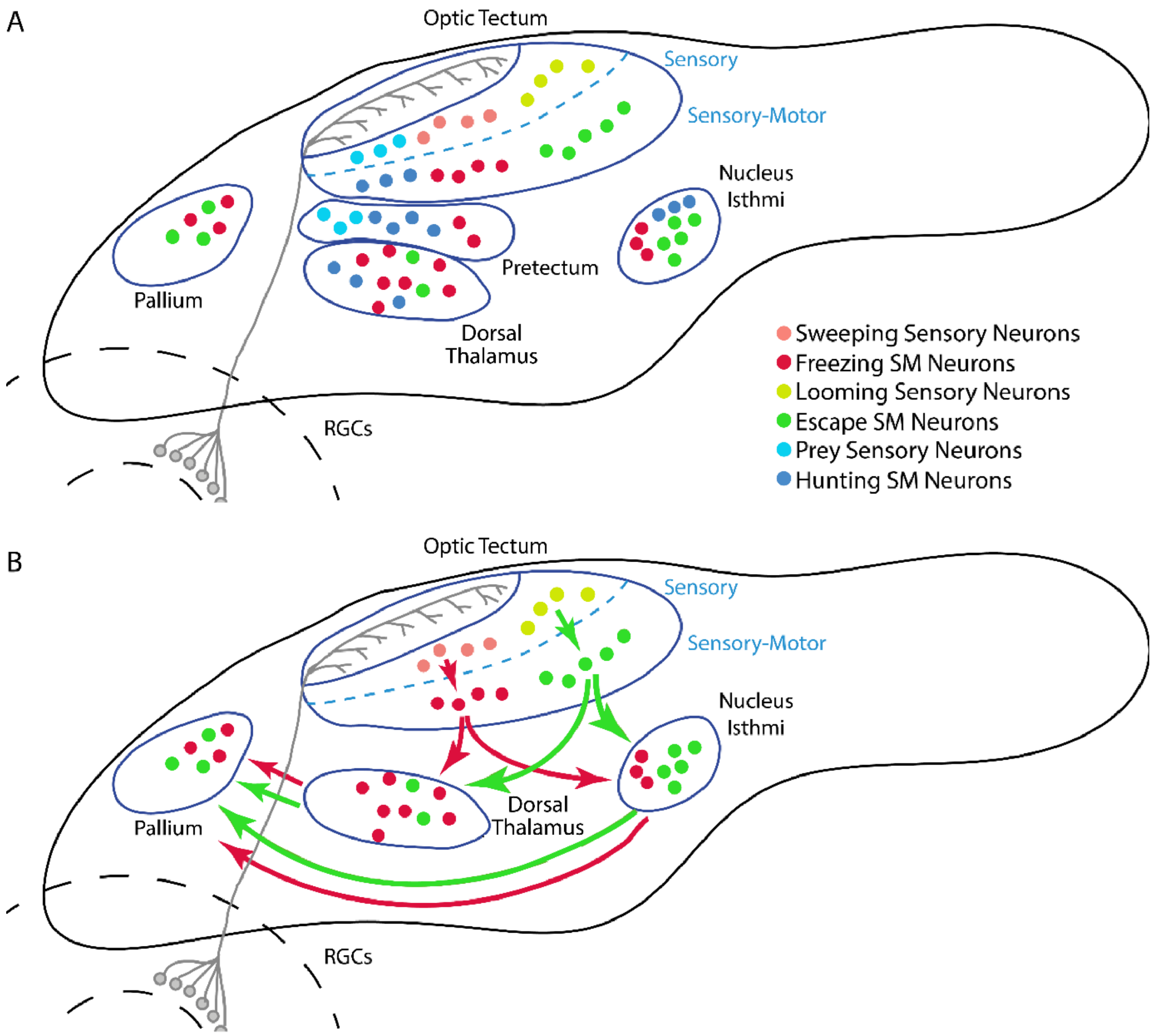


**Supplementary Figure 20: Sensory motor transformation in the visual system of larval zebrafish.** (A) Schematic of the locations of sensory and sensory-motor (SM) neurons in the brain. (B) Proposed defensive behavior pathways based on homogous areas in rodents.

**Supplementary Table 1: Number of Segmented Neurons in Each Brain Region**

| Region | Number of Neurons Segmented |
|---|---|
| All forebrain areas | 47,048 |
| Telencephalon | 25,273 |
| Olfactory Bulb | 3,615 |
| Pallium (dorsal telencephalon) | 18,498 |
| Subpallium (ventral telencephalon) | 3,160 |
| Eminentia Thalami | 1,549 |
| Prethalamus (alar prosomere 3, ventral thalamus) | 1,190 |
| Epiphysis | 78 |
| Habenula | 2,840 |
| Dorsal Habenula | 960 |
| Ventral Habenula | 1,880 |
| Thalamus Proper | 8,970 |
| Pretectum | 13,071 |
| Posterior Tuberculum (basal part of prethalamus and thalamus) | 276 |
| All midbrain areas | 53,160 |
| Tegmentum | 7,926 |
| Tectum | 40,824 |
| Periventricular layer | 32,886 |
| Tectal Neuropil | 7,843 |
| Torus Longitudinalis | 342 |
| Torus Semicircularis | 3,969 |
| All hindbrain areas | 54,036 |
| Cerebellum | 12,862 |
| Medulla oblongata | 41,174 |
| Superior Medulla Oblongata | 32,250 |
| Superior Dorsal Medulla Oblongata | 18,521 |
| Superior Ventral Medulla Oblongata | 13,726 |
| Anterior (dorsal) trigeminal motor nucleus | 977 |
| Posterior (dorsal) trigeminal motor nucleus | 113 |
| Nucleus Isthmi | 10,277 |
| Anterior cholinergic domain of the Nucleus Isthmi | 1,746 |
| Posterior cholinergic domain of the Nucleus Isthmi | 1,258 |

| | |
|---|---|
| Glutamatergic domain of the Nucleus Isthmi | 3,547 |
| Gabanergic domain of the Nucleus Isthmi | 6,287 |
| Superior Ventral Medulla Oblongata (remaining) | 1,494 |
| Locus Coeruleus | 246 |
| Interpeduncular nucleus | 38 |
| Superior Raphe | 584 |
| Intermediate Medulla Oblongata | 8,733 |
| Inferior Medulla Oblongata | 63 |
